# The HLA Ligand Atlas - A resource of natural HLA ligands presented on benign tissues

**DOI:** 10.1101/778944

**Authors:** Ana Marcu, Leon Bichmann, Leon Kuchenbecker, Daniel Johannes Kowalewski, Lena Katharina Freudenmann, Linus Backert, Lena Mühlenbruch, András Szolek, Maren Lübke, Philipp Wagner, Tobias Engler, Sabine Matovina, Jian Wang, Mathias Hauri-Hohl, Roland Martin, Konstantina Kapolou, Juliane Sarah Walz, Julia Velz, Holger Moch, Luca Regli, Manuela Silginer, Michael Weller, Markus W. Löffler, Florian Erhard, Andreas Schlosser, Oliver Kohlbacher, Stefan Stevanović, Hans-Georg Rammensee, Marian Christoph Neidert

## Abstract

The human leukocyte antigen (HLA) complex controls adaptive immunity by presenting defined fractions of the intracellular and extracellular protein content to immune cells. Here, we describe the HLA Ligand Atlas, an extensive collection of mostly matched HLA-I and -II ligandomes from 225 benign samples (29 tissues, 21 subjects). The initial release covers 51 HLA-I and 86 HLA-II allotypes presenting 89,853 HLA-I- and 140,861 HLA-II ligands. We observe that the immunopeptidomes differ considerably between tissues and individuals on both source protein and HLA-ligand level. 1,407 HLA-I ligands stem from non-canonical genomic regions. We highlight the importance of comparatively analyzing both benign and malignant tissues to inform tumor association, based on a case study in three glioblastoma patients. The resource provides insights into applied and basic immune-associated questions in the context of cancer immunotherapy, infection, transplantation, allergy, and autoimmunity. It is publicly available at www.hla-ligand-atlas.org.

## INTRODUCTION

In the past two decades, sequencing the human genome (*genomics*) (Lander et al., 2001; Venter et al., 2001), transcriptome (*transcriptomics*) (Melé et al., 2015), and proteome (*proteomics*) (Kim et al., 2014; Uhlén et al., 2015; Wilhelm et al., 2014) have been milestones that enable a multi-dimensional understanding of biological processes. In the context of the immune system, a subsequent *omics* layer can be defined as the HLA ligandome or the immunopeptidome, comprising the entirety of HLA presented peptides. HLA molecules present peptides on the cell surface for recognition by T cells. These T cells can distinguish self from foreign (Rammensee et al., 1993a, 1993b) peptides, a crucial mechanism in adaptive immunity. Despite HLA-I ligands originating primarily from intracellular proteins, the correlation with their precursors (mRNA transcripts and proteins) is poor (Fortier et al., 2008; Schuster et al., 2017; Weinzierl et al., 2007), limiting approaches based on *in silico* HLA-binding predictions in combination with transcriptomics and proteomics data alone (Boegel et al., 2019; Finotello et al., 2019).

The importance of investigating HLA ligandomes from human healthy and diseased tissues has been well recognized (Caron et al., 2017; Faridi et al., 2018; Vizcaíno et al., 2020) to improve HLA-binding prediction algorithms (Abelin et al., 2019; Bassani-Sternberg et al., 2017; Racle et al., 2019; Reynisson et al., 2020), and immunogenicity prediction analysis (Brown and Holt, 2018; Calis et al., 2013), but also, to inform precision medicine (Freudenmann et al., 2018; Fritsche et al., 2018; Löffler et al., 2019). Direct evidence of naturally presented HLA ligands is required to prove visibility of target peptides to T cells. This is a challenge, for example, in the context of cancer immunotherapy approaches that aim to identify optimal tumor-specific HLA-presented antigens (Bassani-Sternberg et al., 2016; Hilf et al., 2019; Löffler et al., 2019). While their discovery has been made possible by proteogenomics approaches, a major impediment still resides in the lack of benign tissues as a reference for the definition of tumor specificity of target peptides (Chong et al., 2020; Laumont et al., 2018; Schuster et al., 2017). Due to the scarce availability of benign human tissue ligandomes, common alternative strategies are based on transcriptomic datasets either from the same patient, or from multiple tissues extracted from publicly available repositories (Ardlie et al., 2015; Melé et al., 2015). Frequently, morphologically normal tissue adjacent to the tumor (NATs, normal tissues adjacent to tumor) is used as a control in cancer research. However, NATs have been shown to pose unique challenges, since they may be affected by disease and have been suggested to represent a unique intermediate state between healthy and malignant tissues, with a pan-cancer-induced inflammatory response (Aran et al., 2017). Additionally, for some malignancies e.g. of the brain, surgical resection of NATs is inadmissible. Even in cancers that allow for the extraction of NATs, it is still necessary to investigate the presence of potential tumor-associated targets (TAAs) on other tissues to anticipate on-target/off-tumor, systemic adverse events when administering immunotherapies to patients (Cameron et al., 2013; Linette et al., 2013).

In this study we thus employed tissues originating from research autopsies of subjects that have not been diagnosed with any malignancy and have deceased for other reasons, an approach previously described as a surrogate source of benign tissue (Aran et al., 2017; Iacobuzio-Donahue et al., 2019). Although these tissues were affected by a range of non-malignant diseases, we designate their tissues as benign to emphasize morphological normality and absence of malignancy. This definition of benign is in agreement with the definition used by the Genotype-Tissue Expression Consortium (Ardlie et al., 2015; Melé et al., 2015), which provides RNA sequencing data of benign tissues originating from autopsy specimens.

We performed a large-scale mass spectrometry (LC-MS/MS)- based characterization of both HLA-I and -II ligands providing data from benign human tissues obtained at autopsy. The HLA Ligand Atlas is a first draft of a pan-tissue immunopeptidomics reference library from benign tissues comprising for the first time 225 mostly paired HLA-I (198) and -II (217) ligandomes from 29 different benign tissue types obtained from 21 human subjects. For the data analysis, we employed MHCquant (Bichmann et al., 2019), the first open-source customized computational tool for immunopeptidomics assays that provides database search, false discovery rate (FDR) scoring, label-free quantification and binding affinity predictions. In addition, we implemented a user-friendly, web-based interface to query and access the data at https://hla-ligand-atlas.org. Despite its unprecedented comprehensiveness, the HLA Ligand Atlas currently contains only a limited number of tissues and individuals. However, it has been designed as an open and extensible community resource and we strongly encourage the submission of additional data for inclusion. Consistent quality control and data processing will ensure a high quality of the data.

## RESULTS

### The HLA Ligand Atlas: content and scope of the data resource

We describe the HLA Ligand Atlas, a dataset of matched HLA-I and -II ligandomes of benign tissues. HLA-I and -II ligands were isolated via immunoaffinity purification and identified by LC-MS/MS. HLA-binding prediction algorithms and an assessment of peptide length distributions were used to identify high-quality samples and only these were integrated into the dataset (Figure S1 describes the QC steps employed). Our online resource https://hla-ligand-atlas.org provides access to the dataset comprising HLA-I and –II ligands (1% local peptide-level FDR), their source proteins, tissue and subject of origin, as well as all corresponding HLA allotypes classified as strong or weak binders through several user friendly views (Figure 1A, Figure S1). We have acquired HLA ligandome data from 29 distinct tissues obtained from 21 individuals, surmounting to 1,262 LC-MS/MS runs from 225 mostly paired HLA-I (198) and -II (217) samples (Figure 1C, Figure S1, Table S1). The majority of samples was obtained from 14 subjects after autopsy, while 7 additional subjects contributed 5 thymus and 2 ovary samples after surgery. We performed a time series experiment on three benign samples, two ovaries and one liver (Figure S2) and observed no qualitative or quantitative degradation of the immunopeptidome for up to 72 h after tissue removal, supporting the feasibility of employing autopsy tissue as input material for immunopeptidomics assays (Figure S2).

**Figure 1:**
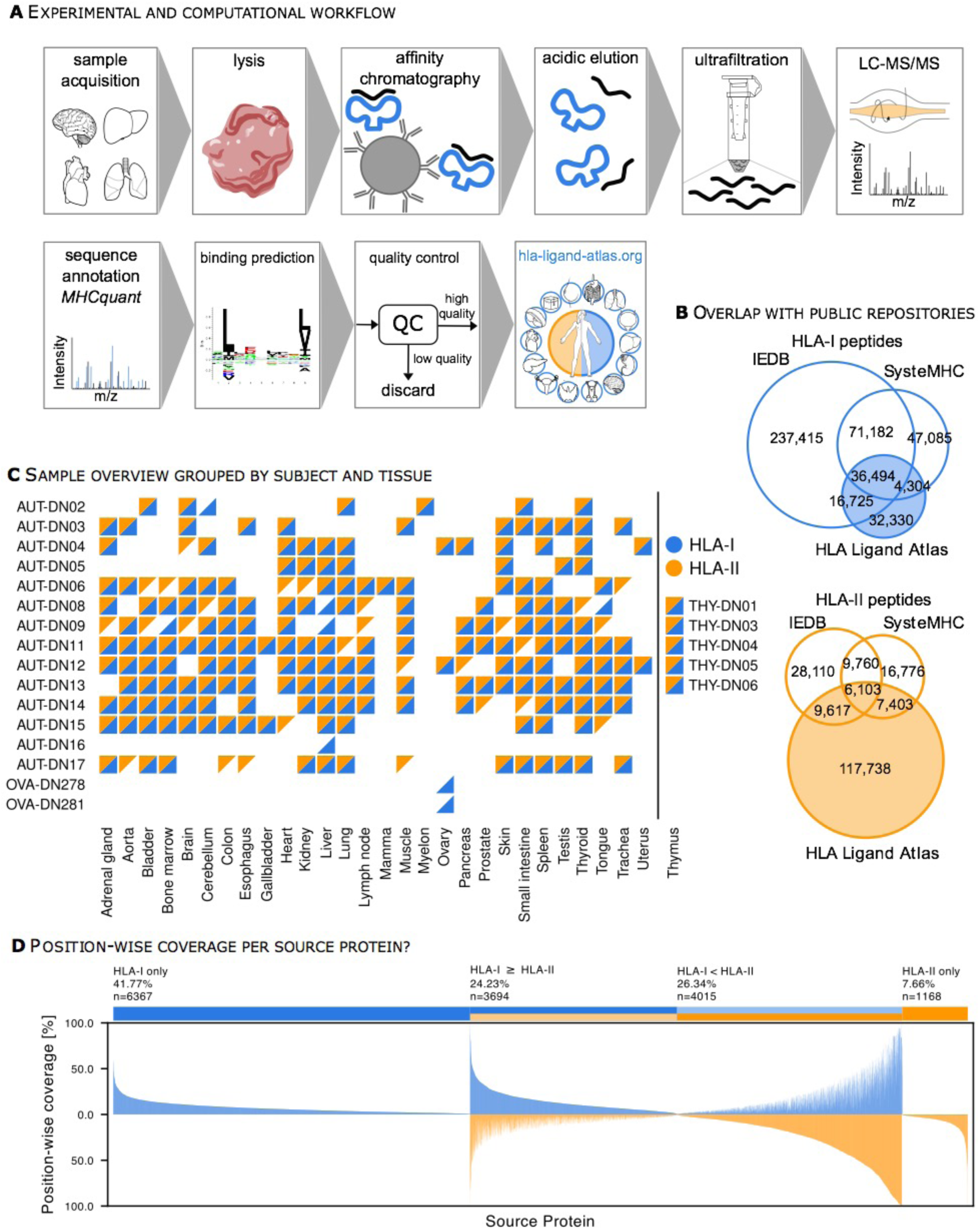
The HLA Ligand Atlas: content and scope of the data resource. (A) The high-throughput experimental and computational workflow steps used to analyze thousands of HLA-I and -II peptides isolated from benign tissues. The resulting HLA-I and -II immunopeptidomes are comprised in the searchable web resource: hla-ligand-atlas.org. See Figure S1 for details of the quality control workflow. See Figure S2 for proof of principle using autopsy tissues. (B) HLA-I and -II peptides expand the know immunopeptidome as curated in the public repositories SysteMHC and IEDB. (C) Sample matrix: HLA-I (blue triangles) and –II samples (orange triangles) included in the HLA Ligand Atlas cover 29 different tissues obtained from 21 human subjects. See Table S1 for patient characteristics. (D) Position-wise coverage (%) of identified source proteins by HLA ligands binned into four groups: (1) exclusively covered by HLA-I peptides, (2) exclusively covered by HLA-II peptides and (3-4) covered by both and separated into higher position-wise coverage by either HLA-I or -II peptides.

Overall, we identified 89,853 HLA-I and 140,861 HLA-II peptides with a local peptide-level FDR of 1% and estimated global peptide-level FDRs of 4.5% and 3.9% for HLA-I and -II peptides, respectively. Identified peptides could be attributed to 51 HLA-I and 81 HLA-II allotypes.

Ultimately, this dataset increases the total number of registered HLA ligands from 413,205 to 445,535 for HLA-I and from 77,769 to 195,507 for HLA-II, as currently encompassed in SysteMHC (Shao et al., 2018) and IEDB (Vita et al., 2015) (Figure 1B).

Moreover, we sought to approximate the worldwide HLA allele frequency of subjects included in the HLA Ligand Atlas. For this purpose, we computed population coverages using the IEDB Analysis Resources (http://tools.iedb.org/population/42) (Table S2). When considering at least one HLA allele match per individual, we observe an allele frequency of 95.1%, 73.6%, 93.0%, for HLA-A (n=16), -B (n=21), and -C (n=14) alleles, respectively. Within the same constraints we observe allele frequencies of 78.8%, 99.5%, 98.2%, 92.3% for HLA-DPB1 (n=9), -DQA1 (n=11), -DQB1 (n=12), and DRB1 (n=19) alleles, respectively (Table S2).

### Source proteins and HLA allotype coverage characteristics of HLA ligands

The HLA Ligands in the dataset were identified based on 15,244 of the 20,365 proteins in Swiss-Prot, hereinafter referred to as source proteins. About half of these source proteins yield both HLA-I and -II ligand identifications, 40% yield only HLA-I ligands and 8% only HLA-II ligands (Figure 1D). We performed a gene ontology enrichment analysis of HLA-I and -II exclusive source proteins, which corroborates the expected cellular compartments associated with the class-specific antigen presentation pathways, with HLA-I presenting primarily intracellular- and HLA-II extracellular proteins (Figure 2F).

**Figure 2:**
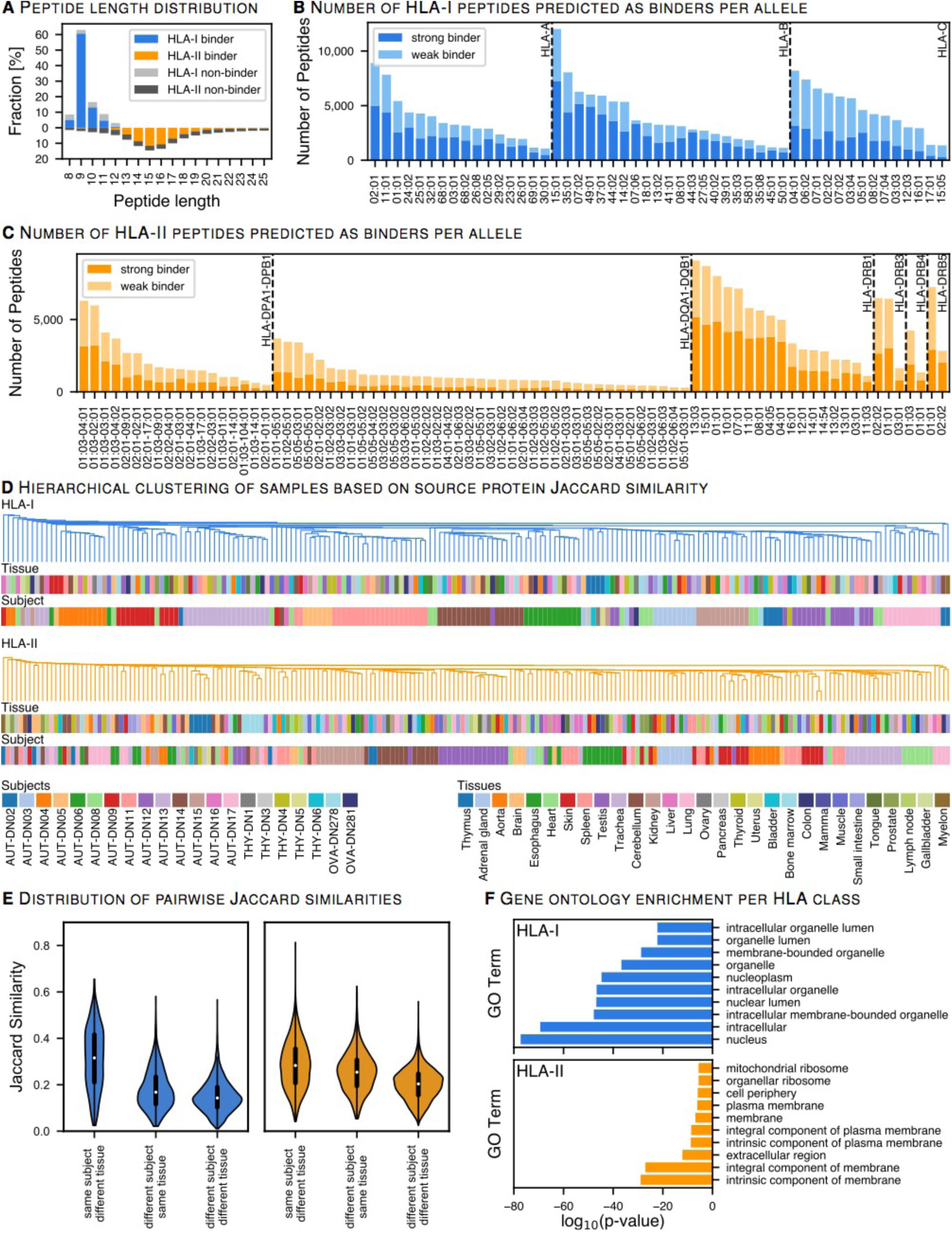
Source proteins and HLA allotype coverage characteristics of HLA ligands. (A) Length distribution of identified HLA-I and -II peptides from all samples was analyzed. HLA-II peptide lengths are mirrored on the negative side of the x-axis. (B, C) Global overview of HLA-I predicted binders distributed across HLA molecules. HLA binding prediction was performed with NetMHCpan 4.0 (% binding rank <2) and SYFPEITHI (Score >50%), while multiple HLA allotypes per peptide were allowed as long as their scores met the aforementioned thresholds. HLA binding prediction for HLA-II ligands was performed with NetMHCIIpan 4.0 and MixMHCPred (% binding rank 0.2 for both) See Figure S3 (D) Pairwise hierarchical clustering of samples based on the Jaccard similarity between HLA-I (blue) and HLA-II (orange) source proteins. The dendrogram illustrates the nearest neighbor based on the similarity between tissues and subjects. See Figure S4 C. (E) Violin plots illustrate the distribution of the Jaccard similarity index for each pairwise comparison between the same subject - different tissues; different subjects - the same tissue, and different subject - different tissues. (F) Gene ontology (GO) term enrichment of cellular components was performed for HLA-I and -II source proteins. Top10 enriched genes with respect to their log10 p-value (Fisher’s exact test) differentiate between intracellular and extracellular antigen processing pathways.

When looking at single amino acid residues across all source proteins (position-wise), 10% of the single residue positions are covered by HLA ligands, a parameter that ranges from 0.02% to 1.9% for individual HLA allotypes (Figure S3). The mode of the overall peptide length distribution depicts the highest abundance of 9mers (60%) for HLA-I and of 15mers (18%) for HLA-II ligands (Figure 2A). While 81% of the HLA-I ligands are predicted to bind a subject’s HLA allotype, this holds true for only 53% of the HLA-II ligands. A major shortcoming of HLA-II binding prediction models appears to be a negative bias towards the tails of the observed peptide length distribution, in particular towards short peptides (Figure 2A). The number of identified peptides that are predicted to bind against specific allotypes varies strongly between allotypes, with HLA-A*02:01, -B*15:01, -B*35:01, -C*04:01 and most HLA-DRB1 allotypes being among the highly represented ones (Figure 2B, C).

### The inter-individual heterogeneity outweighs similarities between tissue types

An unaddressed question, relevant for the discovery and administration of shared TAAs, is if the similarity between tissuetypes outweighs that of individuals. We interrogated the HLA Ligand Atlas and assessed the similarity of the immunopeptidome on both source protein (Figure 2D, E) and HLA-ligand level (Figure S4C, D) between samples, as defined by subject-tissue combinations. For this purpose, we computed pairwise similarities between all samples as measured by the Jaccard similarity index and clustered the samples based on their similarity.

We observe that the sample similarity, even on source protein-level, is dominated by the underlying HLA alleles governing peptide presentation in each subject, resulting in clusters largely reflecting the subjects rather than the tissues. Contrary to our expectations, the five thymus specimens show the same pattern of subject individuality without an increase in source protein overlap with other tissue types. The high subject individuality as indicated by the clustering of subjects rather than tissues holds true irrespective of the data level on which the analysis is based on.

### The immunopeptidome yield varies consistently across tissues

We further investigated the immunopeptidome diversity and variance across all samples for both HLA-I and -II alleles. Overall, we observe a strong variance in the immunopeptidome yield, defined as the number of identified peptides per sample, across all tissues (Figure 3A) and subjects (Figure S4A, B). Despite the inter-individual (i.e., inter-allotype) variance, we can consistently differentiate between high-yielding and low-yielding tissues with respect to both HLA-I and -II peptides (Figure 3, Figure S4A, B). The separation of tissues based on the immunopeptidome yield is not abrupt, but gradual. Low-yielding tissues include skin, aorta, brain, and the gallbladder with a low number of both HLA-I and -II presented peptides across all subjects. On the other hand, high-yielding tissues include thymus, lung, spleen, bone marrow, and kidney (Figure 3A). These tissues have well-characterized immune-related functions or are central components of the lymphatic system. We employed a linear model to systematically evaluate the correlation between the median HLA-I/-II immunopeptidome yield with RNA expression values (RPKM) of immune-related genes identified by targeted RNA sequencing from an external dataset (Boegel et al., 2018) (Figure 3B and Figure S5). We observe a significant correlation between expression values of immune-related genes and HLA-I and -II immunopeptidome yields (Figure S 5A-D). Among these, genes of the immunoproteasome correlate well with the number of HLA-I ligand identifications per tissue (R^2^=0.371, rho=0.669, p=0.002, Figure 3B). Independent studies mapping the healthy human proteome confirm expression of the immunoproteasome in a wide range of tissues, including tissues for which no primary immunological function would be expected (Kim et al., 2014; Wilhelm et al., 2014).

**Figure 3:**
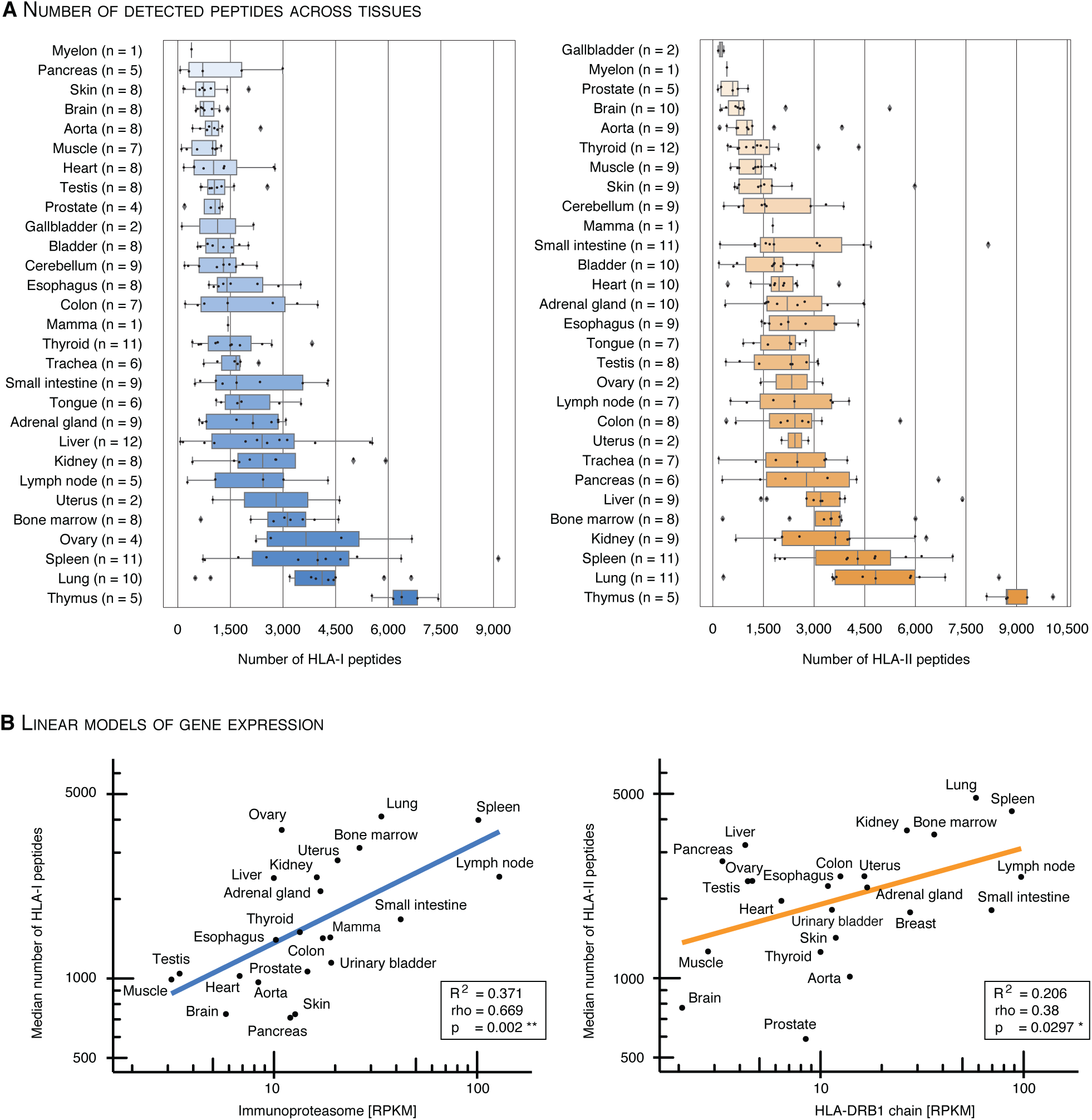
Tissues exhibit a gradual separation based on the immunopeptidome yield. (A) The number of identified HLA-I and -II peptides per sample (subject and tissue combinations) was sorted and plotted by median immunopeptidome yield per tissue. Boxes span the inner two quantiles of the distribution and whiskers extend by the same length outside the box. Remaining outlier samples are indicated as black diamonds. The number of subjects contributing to each tissue is illustrated on the y-axis in parenthesis. (B) A linear model was used to correlate the log transformed HLA-I and -II median peptide yields with log transformed median gene expression counts (RPKM) of the immunoproteasome and HLA-DRB1 per tissue (Boegel et al., 2018). Corresponding R^2^, p-value (F-statistic) and spearman rho are indicated in the bottom right box. See Figure S5

HLA-II peptide yields correlate well with the expression of HLA-DRB1 genes (R^2^=0.206, rho=0.38, p=0.0297, Figure 3B). HLA-DR is well characterized due to the invariant α chain, and thus reduced complexity in the peptide binding groove. Through the high specificity of the L243 antibody for HLA-DR, and the presumably varying specificity of the second antibody Tü39 for different HLA-II allotypes, we cannot exclude a skewed identification in favor of HLA-DRB allotypes. However, higher expression values for HLA-DRB1 compared to other HLA-II allotypes have been described for example in earlier studies on gastric epithelium (Ishii et al., 1992).

### Small subsets of source proteins are tissue-exclusive

Previous studies characterizing the human transcriptome and proteome across tissues have shown varying degrees of tissue-specificity for transcripts and proteins (Jiang et al., 2019; Wang et al., 2019). In this context, we analyzed source proteins of the benign immunopeptidome as a whole and grouped all samples by tissue of origin. We observe a particularly small number of HLA-I (ranging from 5 in mamma to 680 in thymus), and HLA-II (ranging from 8 in ovary to 567 in thymus) source proteins identified exclusively in one tissue (Figure 4A, B, Table S5). Concordantly, only small numbers of tissue-exclusive protein identifications have been observed in human tissue-wide proteomics studies (Wilhelm et al., 2014). Only recently, the systematic, quantitative analysis of the human proteome and transcriptome in multiple tissues has revealed that differences between tissues are rather quantitative than defined by the presence or absence of certain proteins (Jiang et al., 2019; Wang et al., 2019).

**Figure 4:**
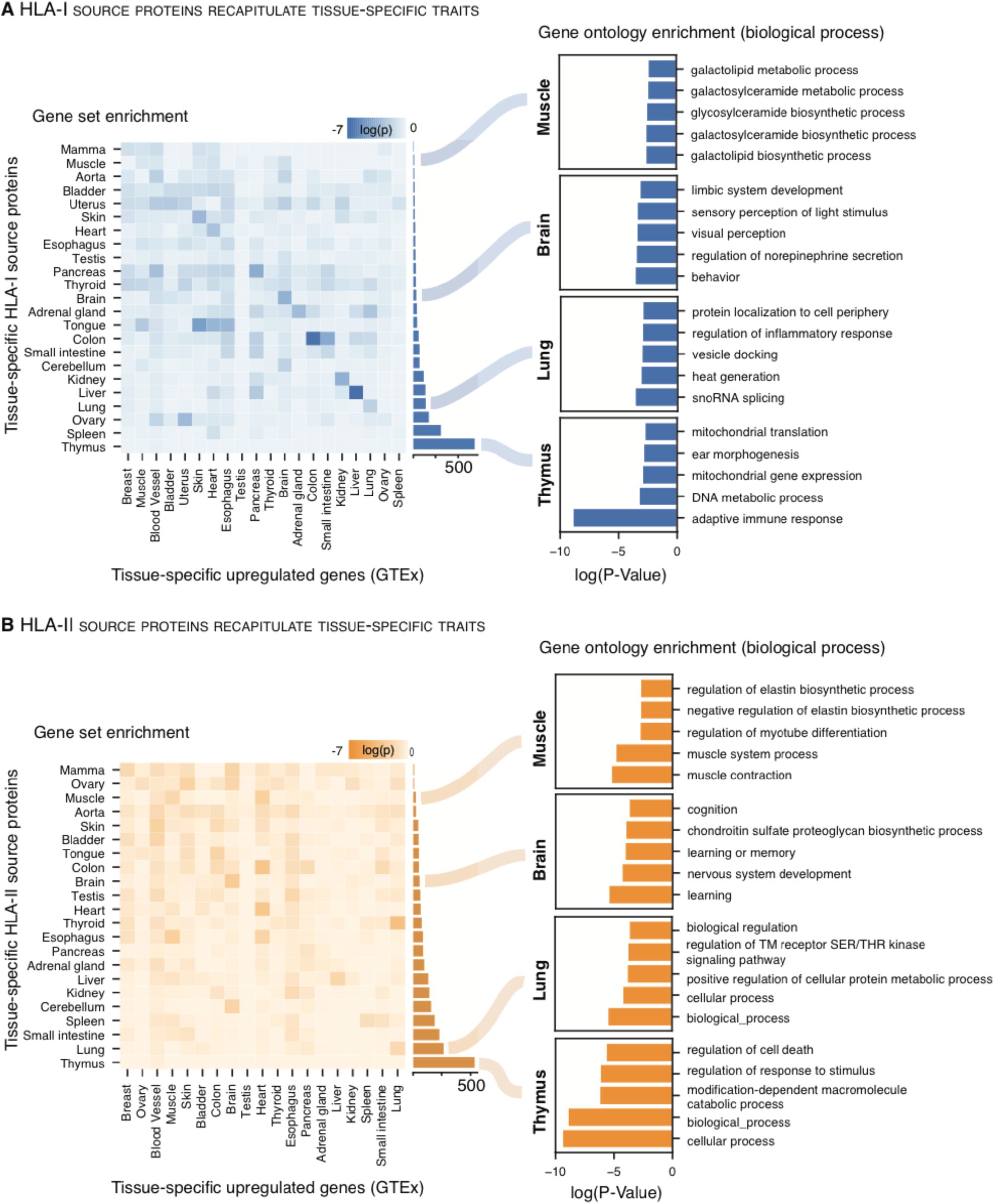
Small subsets of source proteins are tissue exclusive. See Table S5. (A, B) Gene set enrichment (left) was tested for each tissue by correlating unique HLA-I and –II source proteins per tissue with upregulated genes as annotated in GTEx. Heatmaps depict log10 p-values (Fisher’s exact test) for each pairwise comparison. The number of tissue-specific HLA-I and –II source proteins is depicted by the bar plot for each tissue at the right-hand side of the heatmaps. In addition, GO term enrichment (right) of biological processes was performed using the panther DB webservice for selected tissues with the same set of HLA-I and -II tissue-specific source proteins. Top 5 enriched terms with respect to their log10 p-value (Fisher’s exact test) were selected.

Next, we sought to determine whether tissue-specific biology is conserved between the transcriptome and immunopeptidome.

For this purpose, we compared tissue-enriched gene sets from the GTEx repository with tissue-exclusive HLA-I and -II source proteins (Figure 4A, B, left). We observe that tissue-specific biology is represented by HLA-I and -II source proteins through an enrichment with upregulated transcripts in the respective tissue. Gene set enrichment analysis further reflects functional proximity between tissues such as tongue, heart and muscle or brain and cerebellum.

We additionally observed that tissue-specific traits are recapitulated by gene ontology (GO) term enrichment of biological processes (Figure 4A and B, right). Enriched GO terms reveal tissue-specific biological functions such as ‘adaptive immune response’ in the thymus or ‘behavior’ in the brain. However, clear associations between enriched gene sets and HLA-I and -II source proteins are less evident in tissues such as spleen or testis, despite the disparity of tissue-exclusive HLA-I source protein identifications, accounting for only 23 in testis while spleen yields 309.

Overall, tissue-specific traits are more evident for HLA-I than for HLA-II source proteins, as supported by a higher significance, when assessing the correlation between tissue-exclusive source proteins with GTEx-enriched transcripts and function-specific GO terms. HLA-II source proteins are represented by more general GO terms, which still reflect distinct biological processes characteristic for the respective tissue.

### Cryptic peptides are part of the benign immunopeptidome

Recently, cryptic HLA peptides come into focus as a new potential source of tumor-associated antigens (TAAs). Cryptic peptides originate from non-coding regions, i.e. 5’- and 3’-UTR, non-coding RNAs (ncRNA), intronic and intergenic regions, or from shifted reading frames in annotated protein coding regions (off-frame). Ribosome profiling and immunopeptidomics studies confirm their translation and presentation on HLA-I molecules (Chong et al., 2020; Laumont et al., 2018; Ouspenskaia et al., 2020). So far, cryptic peptides have predominantly been characterized in tumors, while their presentation in benign tissues remains poorly charted. We analyzed the HLA-I-restricted LC-MS/MS data of the HLA Ligand Atlas with Peptide-PRISM (Erhard et al., 2020) (Figure 5A) and identified 1,407 cryptic peptides, including the peptide SVASPVTLGK that was classified as a TAA in lung cancer tissue in a previously published study (Figure 5, Table S3) (Laumont et al., 2018). This peptide was identified in the HLA Ligand Atlas in five different subjects in lung and liver tissues. We find that 47% of cryptic peptides were identified in more than one subject (Table S3). Both cryptic and conventional peptides share similar physicochemical properties. Their predicted chromatographic retention time correlates with their experimentally observed retention time equally well as for conventional peptides (Figure 5D) (Chong et al., 2020; Mylonas et al., 2018; Ouspenskaia et al., 2020; Rolfs et al., 2019). The identified cryptic HLA-I ligands can be classified into following genomic categories with decreasing frequency: 5’-UTR (51%), followed by Off-Frame (33%), ncRNAs (13%), 3’-UTR (2%), intergenic (1%), and intronic regions (0.5%) (Figure 5C). The predominance of cryptic peptides from the 5’-UTR is in accordance with previous studies (Erhard et al., 2020; Ouspenskaia et al., 2020). Overall, HLA allotypes show different presentation propensities of cryptic peptides, when related to cryptic and canonical peptides, with HLA-A*03:01 covering the largest fraction of all identified cryptic peptides, followed by -B*07:02 and -A*68:01, as previously observed (Figure 5B) (Erhard et al., 2020).

**Figure 5:**
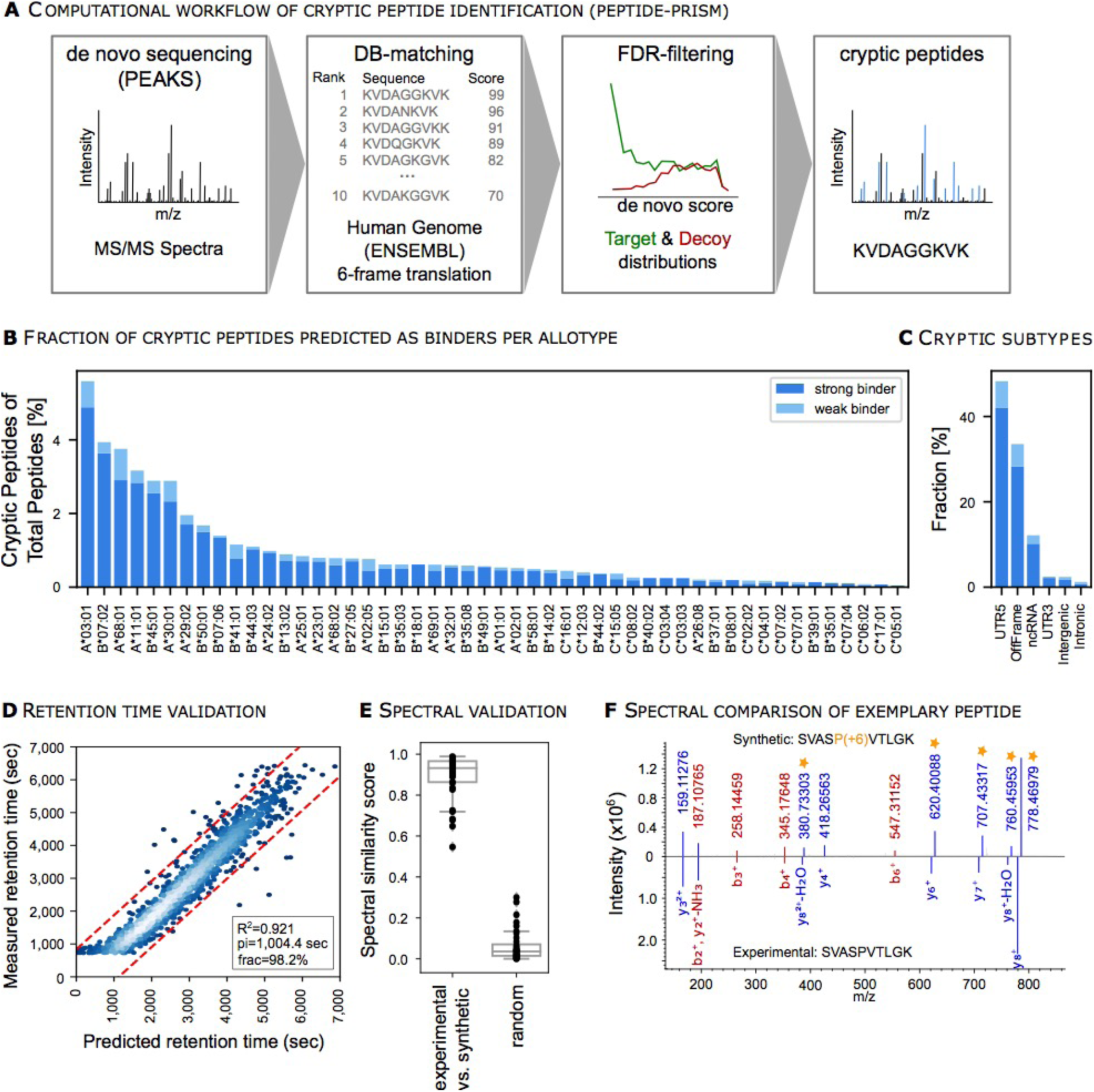
Cryptic peptides are part of the benign immunopeptidomes. See Table S3, Figure S6. (A) Spectra were searched with Peptide-PRISM to identify peptides of cryptic origin. Briefly, *de novo* sequencing was performed, and top 10 sequences per spectra were queried against a database consisting of the 3-frame translated transcriptome (Ensemble 90). Target-Decoy search was performed per database stratum, separately for canonical and cryptic peptides. (B) The HLA-allotype distribution of cryptic peptides was plotted in relation to cryptic and canonical peptides predicted to bind to the respective HLA allotype across all subjects and tissues. (C) Distribution of identified cryptic peptides categorized into multiple non-coding genomic regions. (D) Linear model correlating measured retention times (RT) of cryptic peptides with their predicted RTs trained on canonical peptide RTs. Corresponding R^2^, pi (width of the prediction interval – red dashed lines), and frac (the number of peptides falling into the prediction interval) are indicated in the bottom right. (E) 36 cryptic peptides were selected for spectral validation with synthetic peptides. The similarity between the synthetic and experimental spectrum was computed by correlation scores. F) Exemplary spectral comparison of the cryptic peptide SVASPVTLGK and its synthesized heavy isotope-labeled counterpart (P+6). Matching b (red) and y ions (blue) are indicated as well as the isotope mass shifted ions (orange stars) of the synthesized peptide.

We selected 36 top-ranking (1% FDR) cryptic peptides, shared among subjects for spectral validation by experimental comparison with the corresponding isotopically-labeled synthetic peptide (Table S3). We computed a similarity score between the spectra obtained from the experimental vs. synthetic peptides (Figure 5E, Table S3). A size-matched set of randomly selected comparisons was employed to create a reference negative distribution of the spectral similarity score. We were able to confirm the correct identification of selected cryptic HLA-I ligands, not only based on the computed similarity score, but also through individual inspection (Figure S6). Therefore, we can show that cryptic peptides are not *per-se* tumor-specific, albeit their frequency might be reduced in benign tissues (Erhard et al., 2020)

### HLA Ligand Atlas data enables prioritization of tumor-associated antigens (TAAs)

A general lack of multi-tissue immunopeptidomics reference libraries from benign tissues has been mentioned in previous studies aiming to identify TAAs (Chong et al., 2020; Granados et al., 2016). Here, we propose the implementation of the HLA Ligand Atlas as a reference library of benign multi-tissue immunopeptidomes for comparative profiling with tumor immunopeptidomes for the discovery of actionable TAAs. As a case study, we selected three glioblastoma tumor samples from different individuals and analyzed their immunopeptidomes. We comparatively profiled the HLA-I and -II ligands of the glioblastoma samples against the benign dataset encompassed in the HLA Ligand Atlas (Figure 6A and B). The majority of HLA ligands is shared between both tumor and benign tissues, with 5,185 HLA-I TAAs and 3,246 HLA-II TAAs being unique to glioblastoma (Table S4). When assessing their presentation frequency, 691 HLA-I TAAs are found on two glioblastoma samples, while 4,495 are patient-individual. In the case of HLA-II TAAs, 43 are shared between two glioblastoma patients, and 3,203 are patient-individual. No identified HLA-I or –II ligands were common to all three glioblastoma patients.

**Figure 6.**
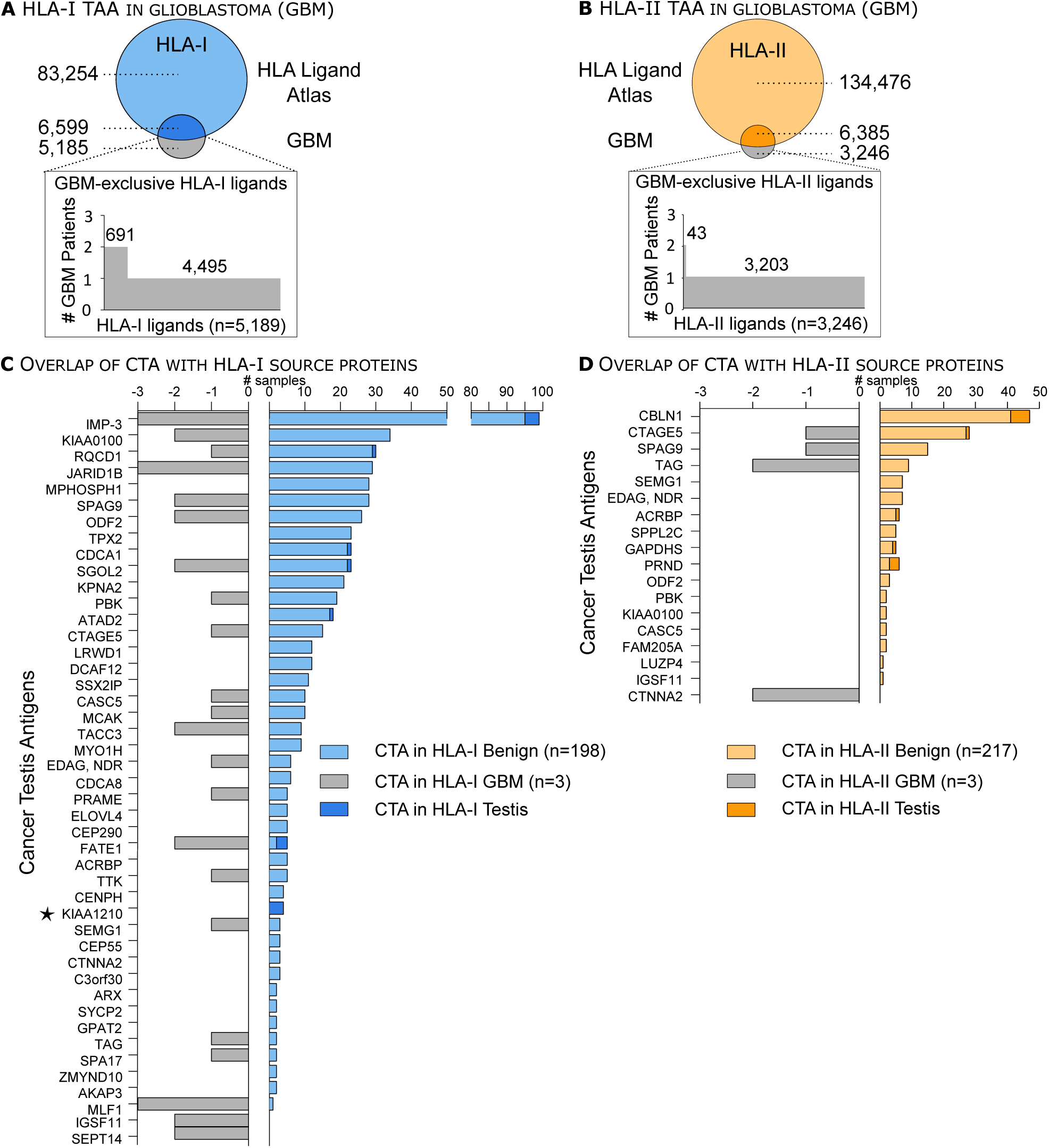
HLA Ligand Atlas data enables prioritization of tumor-associated antigens (TAAs). A, B) The size-proportional Venn diagram illustrates the overlap between the pooled glioblastoma (GBM) and benign HLA-I and -II immunopeptidomes, respectively. The waterfall plots show the number of glioblastoma-associated HLA-I ligands and their frequency among the three glioblastoma (GBM) patients analyzed. See Table S4. C, D) Published CTAs are presented as HLA-I or -II ligands on benign tissues, including testis but also in glioblastoma tumors. The number of identified samples either from the HLA Ligand Atlas or the glioblastoma dataset is depicted on the x-axis, provided that each CTA has been identified with at least two different HLA ligands. The CTA KIA1210 was identified exclusively on HLA-I source proteins in testis and is marked with an asterisk. See Table S4.

Moreover, we investigated the presentation of cancer testis antigens (CTAs) by HLA-I and -II molecules on benign tissues. CTAs are immunogenic proteins preferentially expressed in normal gametogenic tissues and different types of tumors (Almeida et al., 2009; Wang et al., 2016). We compiled a list of 422 published CTAs from the curated CT database (Almeida et al., 2009) and a recent publication aiming to identify CTAs from transcriptomics datasets (Wang et al., 2016). Of 422 published CTAs, 40 CTAs were presented on either HLA-I or -II molecules and 10 CTAs on both HLA-I and -II molecules in the HLA Ligand Atlas, provided that respective source proteins were identified with at least two HLA ligands (Figure 6B, C, Table S3). CTAs, such as IMP-3, KIA0100, and CBLN1 were presented in numerous benign samples with HLA-I and -II ligands (Figure 6C and D, Table S4). Furthermore, the CTA KIA1210 was only identified in the benign dataset on testis in accordance to its CTA status. Similarly, we queried all glioblastoma source proteins against the selected 422 CTAs and found three CTAs (two HLA-I and one HLA-II) exclusively presented in glioblastoma and not in our benign dataset, indicating promising targets against this tumor entity.

### HLA ligands form hotspots in source proteins

When looking at the position-wise coverage profiles of individual source proteins across all HLA allotypes, we observe that HLA ligands seem to emerge from spatially clustered hotspot regions while other areas of the source protein do not contribute any HLA ligands at all (Figure 7, left). It has been shown previously that this clustering effect cannot be explained by the occurrence of HLA binding motifs as incorporated in epitope prediction tools (Müller et al., 2017). The hotspot locations often coincide between HLA-I and -II ligands, however, we did not perform a large-scale statistical analysis to validate this class linkage effect. In the case of HLA-II, the clustering effect has to be distinguished from the co-occurrence of HLA-II ligand length variants, which leads to a large number of distinct peptides covering the same source protein position due to the nature of HLA-II antigen processing and binding (Álvaro-Benito et al., 2018). Many of the observed clusters span ranges of distinct, non-overlapping HLA-II ligands (Figure 7, right). Position-wise coverage plots for all source proteins are available online at hla-ligand-atlas.org.

**Figure 7:**
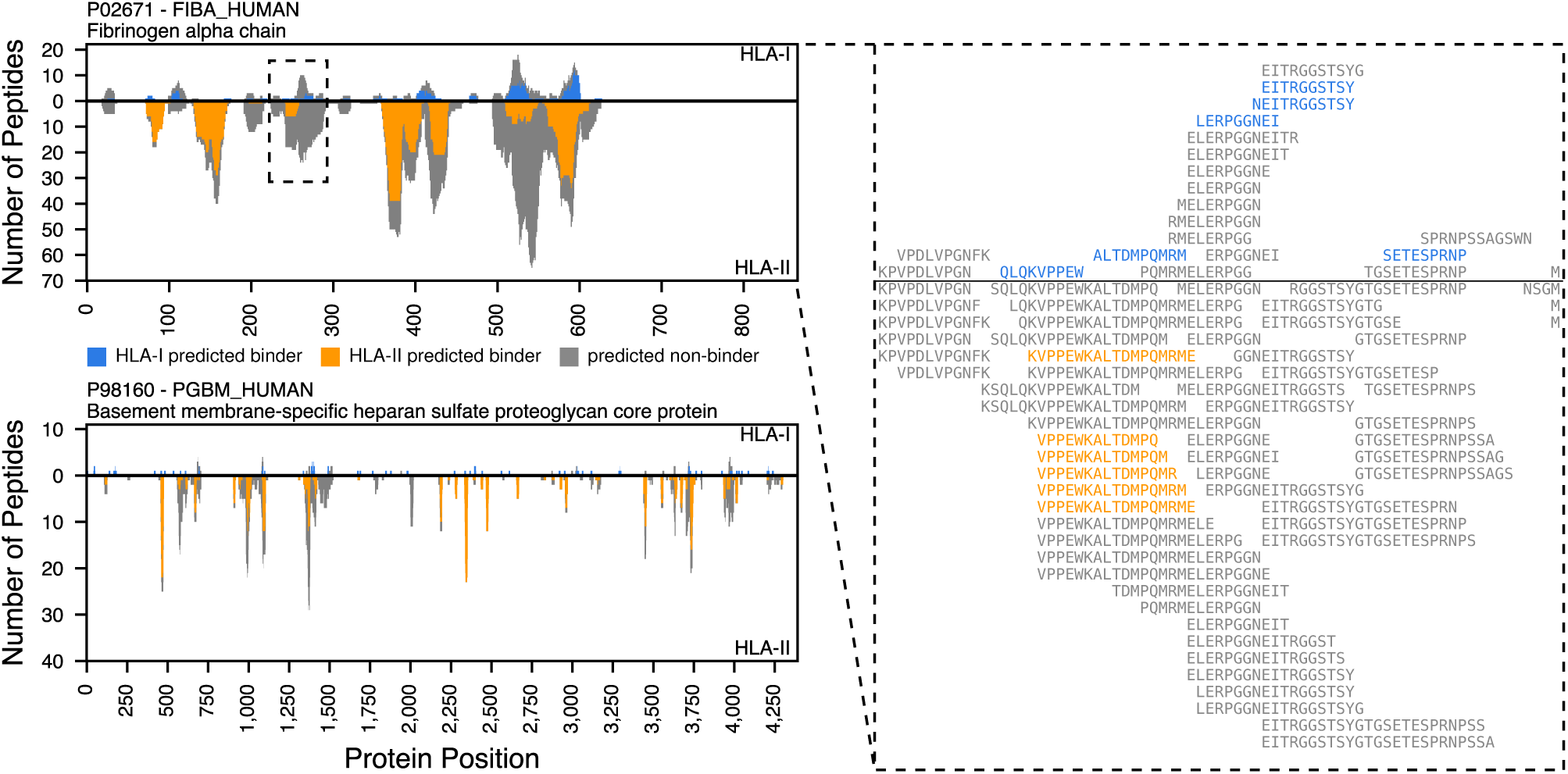
HLA ligands form hotspots in source proteins. The position-wise HLA ligand coverage profiles as available in the HLA Ligand Atlas web interface for two exemplary proteins (left), the fibrinogen alpha chain (Uniprot ID P02671, length 866 aa, top) and the basement membrane-specific heparan sulfate proteoglycan core protein (Uniprot ID P98160, length 4,391 aa, bottom) are shown, illustrating the spatial clustering of HLA ligands into hotspots. For P02671 a close-up of such a cluster is shown in form of a multiple sequence alignment of the identified peptides (right).

### The HLA Ligand Atlas web interface

The HLA Ligand Atlas web interface was designed to allow users to conveniently access the data we collected. Users can formulate queries in the form of filters based on peptide sequences, peptide sequence patterns, HLA allotypes, tissues and proteins of origin, or combinations thereof. Additionally, users can submit files with peptides or uniport IDs, either as plain lists or as a FASTA files. The peptide list is then queried against the database and the resulting hits can again be freely combined with the aforementioned filters. Query results are shown as a list of peptides with plots of the corresponding HLA allotype and tissue distributions. Additionally, detailed views for single peptides and for coverage of proteins are available. Apart from the query interface, the web front-end also displays various aggregate views of the data stored in the database.

## DISCUSSION

In this study, we create a novel data resource termed the HLA Ligand Atlas which is publicly available and easily searchable at hla-ligand-atlas.org (Figure 1C). It provides for the first time a comprehensive collection of benign human HLA-I and -II immunopeptidomes. The large number of different HLA allotypes will help to considerably improve HLA-binding prediction algorithms, particularly for infrequent HLA alleles. HLA-II immunopeptidomes paired with high-resolution HLA-II typing are still scarcely available and therefore represent a valuable resource for improving HLA-II prediction models.

We find that HLA allotypes display varying presentation propensities towards certain peptide populations, with HLA-B*15:01 and HLA-DRB1*01:01 presenting the highest number of canonical self-peptides, and HLA-A*03:01 and B*07:02 presenting the highest proportion of cryptic peptides in our subject cohort and a previously published study (Erhard et al., 2020). The increased number of peptides presented on a subset of HLA alleles can be attributed to their frequency among the analyzed individuals or to their potentially high copy number on cells. Further technical biases can influence the immunopeptidome yield, such as antibody preferences towards certain HLA allotypes, ionization and fragmentation characteristics of eluted HLA ligands, but also binding prediction algorithms that perform better for frequent, well studied HLA allotypes. However, HLA allotypes have evolved to present different peptide subsets to T cells (Kaufman, 2018), examples ranging from HLA-B*40 being able to stabilize the negative charge of phosphorylated peptides (Alpízar et al., 2017), and HLA-B*57 conferring a survival advantage in HIV infections (Marino, 2018; Vizcaíno et al., 2020). Moreover, we observed multiple HLA allele matches per peptide which is indicative of binding similarities between HLA allotypes, or promiscuous HLA alleles that allow binding of a large repertoire of different peptides (Kaufman, 2018).

One fundamental and so far unanswered question concerns the similarity of immunopeptidomes across individuals. Our evaluation of the Jaccard similarity index across samples in the HLA Ligand Atlas provides evidence that differences between individuals exceed differences between tissue types in the same individual for both the immunopeptidome and their source proteins. On a proteome level, however, samples were previously separated by tissue type, rather than individuals (Jiang et al., 2019). Nonetheless, HLA-allotype-dependent selection and editing throughout the antigen presentation pathway shape the immunopeptidome, complicating its prediction from genomic, transcriptomic, and proteomic data sources. While we analyzed 21 human subjects in this study, a larger number would be required to answer this question unequivocally.

The high degree of individuality between immunopeptidomes, even when subjects share a subset of HLA allotypes, has major repercussions for clinical applications in emerging fields such as immuno-oncology. Our findings indicate that the immunopeptidome adds an additional layer of complexity to the well-described genomic and transcriptomic tumor-heterogeneity. Successful induction of T cell responses after peptide vaccination with neoantigens (Ott et al., 2017; Sahin et al., 2017) indicate that precision medicine will evolve to an increasingly individualized field, where treatment options will be tailored to the immunopeptidomic landscape of the tumor. Mapping the tumor HLA ligandome of an individual patient therefore needs to be paralleled by a broad and in-depth knowledge of its benign counterpart – the HLA Ligand Atlas is a first step in this direction. When dissecting the HLA Ligand Atlas tissue-wise, we observe a paired immunopeptidome yield between HLA-I and -II ligands that could be indicative of an increased infiltration of immune cells in high-yielding tissues. Alternatively, expression of HLA-II molecules on cells other than APCs could explain this observation. By analyzing bulk tissue, a definite statement whether peptide presentation occurred on tissue or tissue-infiltrating immune cells cannot be made. The immunopeptidome yields per tissue correlate positively with RNA expression profiles of genes related to antigen processing and presentation. Yet, the identified source proteins appear to be barely specific for the tissue of origin. The weak correlation between immunopeptidome yield and RNA expression values has been observed previously (Fortier et al., 2008; Schuster et al., 2017). Although abundant HLA ligands stem from highly expressed transcripts, most HLA ligands span a wider dynamic range of gene expression (Wang et al., 2019). Furthermore, it was recently shown that the immunopeptidome is better captured by the translatome as identified by ribosome profiling than by the transcriptome (Chong et al., 2020; Ouspenskaia et al., 2020).

Low-yielding samples barely display any tissue-specific proteins. However, tissue-specific source proteins often reflect tissue-specific traits when correlated to enriched gene sets of GTEx transcriptomes from the respective tissues. Therefore, tissue-specific function is represented in the immunopeptidome, but differences between tissues cannot be imputed from the immunopeptidome alone. Studies mapping the whole proteome in multiple human tissues report few proteins with tissue-specific expression (Wang et al., 2019; Wilhelm et al., 2014) and suggest that differences between tissues might be quantitative, and less dominated by the presence or absence of protein species (Jiang et al., 2019; Wang et al., 2019).

Recent studies have focused on HLA-presented peptides derived from non-coding regions. Ribosome profiling, RNA sequencing, and immunopeptidomics studies have confirmed that cryptic HLA-I peptides expand the known HLA-I immunopeptidome by up to 15% (Erhard et al., 2020), up to 3.3% (Ouspenskaia et al., 2020), and about 10% (Laumont et al., 2018). These studies have mainly focused on tumors and tumor cell lines, PBMCs and mTEC cell lines, in most cases treated and expanded *in vitro*. We employed Peptide-PRISM and identified 1,407 cryptic HLA-I ligands from benign, primary, human samples. Corroborating other studies, we find that a large proportion (41%) are also shared between multiple subjects (Chong et al., 2020; Ouspenskaia et al., 2020).

An essential application of the HLA Ligand Atlas is the selection of candidates for immunotherapy approaches. We propose a workflow to prioritize the large candidate pool of non-mutated tumor-associated targets by comparatively profiling immunopeptidomes of primary tumors and benign tissues, as comprised in the HLA Ligand Atlas. This approach would complement current strategies based on transcriptomes of benign tissues as comprised in GTEx. The HLA Ligand Atlas represents a first draft of a tissue-wide immunopeptidomics map covering both HLA-I and -II canonical peptides, but also HLA-I non-canonical peptides, that can be employed as an orthogonal level of quality control when defining TAAs. Furthermore, anecdotal observations of position-wise coverage of source proteins confirm a previously stated hypothesis, that immunopeptidomes cluster into hotspots of antigen presentation (Bilich et al., 2019; Müller et al., 2017). We envision that different types of TAAs such as mutated, non-mutated, post-translationally modified, of cryptic or proteasomally spliced origin might cluster as well within these hotspots. Future studies that aim to enhance our understanding of such mechanistic patterns of peptide presentation will benefit greatly from the data the HLA Ligand Atlas comprises.

A series of systematic technical limitations in LC-MS/MS-based studies influences the identification depth in each sample. Such aspects include the still limited sensitivity and dynamic range of detection, the insufficient coverage of amino acids in peptide mass spectra, but also shortcomings in peptide identification algorithms. Advances in LC-MS/MS technology, data acquisition methods and computational tools are constantly improving the depth of investigation in immunopeptidomics experiments. Therefore, we encourage the reanalysis of the raw LC-MS/MS dataset with novel hypotheses and upcoming computational methods that will lead to additional insight. Overall, we anticipate that the number of charted human immunopeptidomes will increase, similarly as the human genome and transcriptome were mapped across multiple individuals. By generating larger datasets from many human individuals, population-wide conclusions can be drawn, and immunopeptidome-wide studies will provide insight into disease-associated HLA alleles and peptides (Vizcaíno et al., 2020). The HLA Ligand Atlas strives to advance our understanding of fundamental aspects of immunology relating to autoimmunity, infection, transplantation, cancer immunotherapy and might provide a foundation for vaccine design. We hope that together with the scientific community we can expand the benign immunopeptidome to encompass more human subjects, tissues and HLA alleles.

## ACKNOWLEDGEMENTS

We thank Claudia Falkenburger, Ulrich Wulle, Patricia Hrstic, Nicole Bauer, and Beate Pömmerl for excellent technical support. We also thank Dogukan Yaser, Tobias Kaster, and Jens Bauer for experimental help with the immunopeptidomics experiments. We thank Tjeerd Dijskstra, Etienne Caron, Andreas Friedrich, Carolina Barra, Morten Nielsen, and Hannes Röst for valuable discussions.

This work was funded by the Deutsche Forschungsgemeinschaft (DFG, German Research Foundation) under Germany’s Excellence Strategy - EXC 2180 – 390900677; the Deutsche Forschungsgemeinschaft (DFG) SFB 685 "Immunotherapy: Molecular Basis and Clinical Application“; the ERC AdG 339842 MUTAEDITING; the Boehringer Ingelheim Foundation for Basic Research in Medicine, the Bosch Research Foundation, and Deutsche Forschungsgemeinschaft (DFG) as part of the German Network for Bioinformatics Infrastructure (de.NBI).

## AUTHOR CONTRIBUTIONS

Conceptualization: A.M., L.K., L.Bi., M.C.N., D.J.K., L.Ba., M.W.L.

Methodology: L.Bi., L.K., A.M., D.J.K., L.Ba.

Software: L.Bi, L.K., A.Sz., F.E, A.Sc.

Validation: A.M., L.K., L.Bi., L.K.F.

Formal Analysis: L.Bi., L.K., A.M., A.Sc., F.E., A.Sz., L.K.F.

Investigation: A.M., L.K.F., L.M.

Resources: M.C.N., H-G.R., S.S., O.K., M.W., L.R., H.M., R.M., S.M., T.E., P.W., M.W.L., J.W., M.H-H.

Data Curation.: L.K., A.M., L.Bi., M.C.N., J.V., M.S., K.K.

Writing – Original Draft: A.M., L.K., L.Bi.

Writing – Review & Editing: A.M., L.K., L.Bi., M.C.N., M.W.L., L.K.F., M.L., O.K., S.S., H-G.R., M.W., L.R., H.M., R.M., S.M.,

T.E., P.W., K.K., J.S.W., J.V., M.S., L.Ba., D.J.K., L.M., J.W., M.H-H.

Visualization: L.Bi, L.K., A.M.

Funding Acquisition: H-G.R., S.S., M.C.N., O.K., J.S.W., M.W.L.

## METHODS

### Experimental model and subject details

Human tissue samples were obtained *post-mortem* during autopsy performed for medical reasons at the University Hospital Zürich. The study was approved by the Cantonal Ethics Committee Zürich (KEK) (BASEC-Nr. Req-2016-00604). For none of the included patients a refusal of *post-mortem* contribution to medical research was documented and study procedures are in accordance with applicable Swiss law for research on humans (Bundesgesetz über die Forschung am Menschen, Art. 38). In addition, the study protocol was reviewed by the ethics committee at the University of Tübingen and received a favorable assessment without any objections to the study conduct (Project Nr. 364/2017BO2).

None of the subjects included in this study was diagnosed with any malignant disease. Tissue samples were collected during autopsy, which was performed within 72 hours after death. Tissue annotation was performed by a board-certified pathologist. Tissue samples were immediately snap-frozen in liquid nitrogen.

Thymus samples were obtained from the University Children’ s Hospital Zürich/ Switzerland. Thymus tissue was removed during heart surgery or for other medical reasons. Tissue samples from residual material not required for diagnostic or other medical purposes were obtained after informed consent from the parents of the respective children, in accordance with the principles of the Declaration of Helsinki. The study was approved by the Cantonal Ethics Committee Zürich (KEK) (EC-Nr. 2014-0699, PB_2017-00631) on February 27^th^ 2015.

Furthermore, two benign ovarian tissue samples were collected for the project (OVA-DN278 and OVA-DN281). Both patients were postmenopausal and had a bilateral ovarectomy for cystadenofibromas, which were diagnosed by histopathologic examination of the specimen. The samples were obtained from a normal part of the ovary. The study was approved by the ethical committee of the University of Tübingen (354/2011BO2). Finally, we included three primary glioblastoma tumor samples to illustrate a selection strategy for tumor associated antigens. The primary glioblastoma tumor was removed for patients GBM616 and GBM654, whereas, a recurrent tumor was analyzed for GBM753. The study was approved by the Cantonal Ethics Committee Zürich (KEK) (EC-Nr. 2014-0699, PB_2017-00631).

### HLA typing

Multiple HLA typing approaches were performed for the different sources of patient material.

Autopsy subject AUT-DN08, AUT-DN16, and two benign ovary samples (OVA-DN278 and OVA-DN281) were typed at the Department of Transfusion Medicine of the University Hospital of Tübingen. High-resolution HLA typing was performed by next-generation sequencing on a GS Junior Sequencer using the GS GType HLA Primer Sets (both Roche, Basel, Switzerland). HLA typing was successful for HLA-A, -B, and -C alleles. However, HLA-II typing was only reliable for the HLA-DR locus, and incomplete for the HLA-DP and -DQ loci.

Therefore, we performed exome sequencing of lung tissue for remaining autopsy subjects. Exome sequencing data was processed and OptiType (Szolek et al., 2014) was employed to identify HLA I and -II alleles.

Finally, sequence-based typing was performed for the five thymus samples and the three glioblastoma samples, by sequencing exons 1-8 for HLA-I alleles and exons 2-6 for HLA-II alleles (Histogenetics, Ossining, NY).

The subject characteristics are summarized in Supplemental Table S1 encompassing information on sex, age, the number of collected tissues and HLA-I and II alleles.

### HLA immunoaffinity purification

HLA-I and -II molecules were isolated from snap-frozen tissue using standard immunoaffinity chromatography. The antibodies employed were the pan-HLA-I-specific antibody W6/32 (Barnstable et al., 1978), and the HLA-DR-specific antibody L243 (Goldman et al., 1982), produced in house (University of Tübingen, Department of Immunology) from HB-95, and HB-55 cells (ATCC, Manassas, VA) respectively. Furthermore, the pan-HLA-II-specific antibody Tü39 was employed and produced in house from a hybridoma clone as previously described (Pawelec et al., 1985). The antibodies were cross-linked to CNBr-activated sepharose (Sigma-Aldrich, St. Louis, MO) at a ratio of 40 mg sepharose to 1 mg antibody for 1 g tissue with 0.5 M NaCl, 0.1 M NaHCO3 at pH 8.3. Free activated CNBr reaction sites were blocked with 0.2 M glycine.

For the purification of HLA-peptide complexes, tissue was minced with a scalpel and further homogenized with the Potter-Elvehjem instrument (VWR, Darmstadt, Germany). The amount of tissue employed for each purification is documented in Supplemental Table S1. This information is not available for seven tissues, annotated as n.d. in said table. Tissue homogenization was performed in lysis buffer consisting of CHAPS (Panreac AppliChem, Darmstadt, Germany), and one cOmplete™ protease inhibitor cocktail tablet (Roche) in PBS. Thereafter, the lysate was sonicated and cleared by centrifugation for 45 min at 4,000 rpm, interspaced by 1 h incubation periods on a shaker at 4°C. Lysates were further cleared by sterile filtration employing a 5 µm filter unit (Merck Millipore, Darmstadt, Germany). The first column contained 1 mg of W6/32 antibody coupled to sepharose, whereas the second column contained equal amounts of Tü39 and L243 antibody coupled to sepharose. Finally, the lysates were passed through two columns cyclically overnight at 4°C. Affinity columns were then washed for 30 minutes with PBS and for 1 h with water. Elution of peptides was achieved by incubating four times successively with 100 – 200 µl 0.2% TFA on a shaker. All eluted fractions were subsequently pooled. Peptides were separated from the HLA molecule remnants by ultrafiltration employing 3 kDa and 10 kDa Amicon filter units (Merck Millipore) for HLA-I and HLA-II, respectively. The eluate volume was then reduced to approximately 50 µl by lyophilization or vacuum centrifugation. Finally, the reduced peptide solution was purified five times using ZipTip Pipette Tips with C18 resin and 0.6 µl bed volume (Merck,) and eluted in 32.5% ACN/0.2% TFA. The purified peptide solution was concentrated by vacuum centrifugation and supplemented with 1% ACN/0.05% TFA and stored at -80°C until LC-MS/MS analysis.

### Time series experiments

We performed time series experiments to assess the suitability of tissues obtained from autopsies as a source of human tissues for the characterization of the benign immunopeptidome. We evaluated the degradation profile of the immunopeptidome, when tissues were stored at 4°C for up to 72 h after tissue removal, to mimic the conditions at autopsy. The time series experiment was repeated in three benign tissues from different individuals: one benign liver obtained at autopsy (AUT-DN16 Liver), and two benign ovaries removed surgically (OVA-DN278 and OVA-DN281). The tissues were extracted and incubated at 4°C until a certain time point and flash-frozen in liquid nitrogen until HLA ligand extraction. As more tissue was available form AUT-DN16 Liver, tissue samples were frozen after 8 h, 16 h, 24 h, 48 h, and 72 h. Due to the limited sample amount obtained from OVA-DN278 and OVA-DN281, only three time points could be accounted for: 0 h, 24 h, and 72 h. The HLA immunoaffinity purification was performed as mentioned, with the exception that mass to volume ratio in ovary samples was adjusted to the lowest mass across all time points before loading onto sepharose columns.

### Mass spectrometric data acquisition

HLA ligand characterization was performed on an Orbitrap Fusion Lumos mass spectrometer (Thermo Fisher Scientific, San Jose, CA) equipped with a Nanospray Flex™ Ion Source (Thermo Fisher Scientific) coupled to an Ultimate 3000 RSLC Nano UHPLC System (Thermo Fisher Scientific). Peptide samples were loaded with 1% ACN/ 0.05% TFA on a 75 µm x 2 cm Acclaim™ PepMap™100 C18 Nanotrap column (Thermo Fisher Scientific) at a flow rate of 4 µl/min for 10 minutes. Separation was performed on a 50 µm x 25 cm PepMap RSLC C18 (Thermo Fisher Scientific) column, with a particle size of 2 µm. Samples were eluted with a linear gradient from 3% to 40% solvent B (80% / 0.15% FA in water) at a flow rate of 0.3 µl/min over 90 minutes. The column was subsequently washed by increasing to 95% B within 1 minute, and maintaining the gradient for 5 minutes, followed by reduction to 3% B and equilibration for 23 minutes.

Data acquisition was performed as technical triplicates in data-dependent mode, with customized top speed (3 s) methods for HLA-I- and HLA-II-eluted peptides. HLA-I peptides have a length of 8 - 12 amino acids (Rammensee, 1995; Stern et al., 1994), therefore, the scan range was restricted to 400 - 650 m/z and charge states of 2 - 3. MS1 and MS2 spectra were detected in the Orbitrap with a resolution of 120,000 and 30,000 respectively. Furthermore, we set the automatic gain control (AGC) targets to 1.5*105 and 7.0*104 and the maximum injection time to 50 ms and 150 ms for MS1 and MS2, respectively. The dynamic exclusion was set to 7 s. Peptides were fragmented with collision-induced dissociation (CID) while the collision energy was set to 35%.

HLA-II peptides have a length of 8 - 25 amino acids (Lippolis et al., 2002; Stern et al., 1994), thus the scan range was set to 400 -1,000 m/z and the charge states were restricted to 2 -5. Readout for both MS1 and MS2 were performed in the Orbitrap with the same resolution and maximum injection times as for HLA-I peptides. The dynamic exclusion was set to 10 s and AGC values employed were 5.0*105 and 7.0*104 for MS1 and MS2, respectively. Higher-energy collisional dissociation (HCD) fragmentation with 30% collision energy was employed for HLA-II peptides.

### Database search with MHCquant

MS data obtained from HLA ligand extracts was analyzed using the nf-core (Ewels et al., 2020) containerized, computational pipeline MHCquant (Bichmann et al., 2019) (release 1.5.1 - https://www.openms.de/mhcquant/) with default settings. The workflow comprises tools to analyze LC-MS/MS data of the open-source software library OpenMS (2.5) (Röst et al., 2016). Identification and post-scoring were performed using the OpenMS adapters to Comet 2016.01 rev. 3 (Eng et al., 2015) and Percolator 3.4 (The et al., 2016) at a local peptide-level false discovery rate (FDR) threshold of 1% among the technical replicates per sample. Subsequently, we estimated the global peptide-level FDR by dividing the sum of expected false positive identifications from each sample (1% peptide level FDR) by the total number of identified peptides in the entire dataset (HLA-I: 4.5% FDR, HLA-II: 3.9% FDR) (Reiter et al., 2009; Savitski et al., 2015). The human reference proteome (Swiss-Prot, Proteome ID UP000005640, 20,365 protein sequences) was used as a database reference. Database search was performed without enzymatic restriction, with methionine oxidation as the only variable modification. MHCquant settings for high-resolution instruments involving a precursor mass tolerance of 5 ppm and a fragment bin tolerance of 0.02 Da were applied. The peptide length restriction, digest mass and charge state range were set to 8-12 amino acids, 800-2500 Da and 2-3 for HLA-I and 8-25 amino acids, 800-5000 Da and 2-5 for HLA-II, respectively.

### HLA binding prediction

Peptide binding predictions were computed based on the subject’s HLA alleles. For HLA-I ligand extracts, we employed SYFPEITHI (Rammensee et al., 1999) and NetMHCpan-4.0 (Jurtz et al., 2017) in ligand mode (default). The SYFPEITHI score *S*_SyF_ was computed by dividing the sum of amino acid-specific values for each position in the tested peptide by the maximally attainable score for the respective HLA allotype (Di Marco et al., 2017).

HLA-II ligand extracts were annotated with NetMHCIIpan-4.0 (Reynisson et al., 2020) and MixMHC2pred (Racle et al., 2019) using the default settings.

Peptides were categorized as strong binders against a given HLA allotype if either netMHCpan-4.0, netMHCIIpan-4.0 or MixMHC2pred reported a percentile rank score *S*_rank_ ≤.0.5. Peptides were reported as weak binders if any of the tools reported *S*_rank_ ≤ 2.0 or in case of SYFPEITHI *S*_SYF_ ≤ 0.5 All peptide-HLA allotype associations within these limits were included in the dataset, i.e., a single peptide sequence can be reported as a binder against multiple allotypes of the same donor. Unless allele associations are specified, all peptides including classified non-binders against any subject’s allotype were included in the analysis.

### Binding prediction and length distribution-based quality control

We defined the fraction of predicted binders of a sample as the ratio of predicted binders divided by the total number of peptide identifications. Technical replicates with a fraction of predicted binders lower than 50% for HLA-I and lower than 10% for HLA-II ligand extracts were excluded from the dataset. Furthermore, individual replicates were removed from the dataset if the mode of the length distribution differed from 9 amino acids for HLA-I and was not in the interval [12, 18] for HLA-II (see Figure S1).

### Quantitative time series analysis

Database search of LC-MS/MS data from the three time series experiments was performed with MHCquant 1.5.1 as previously described (Bichmann et al., 2019). Identifications were matched between runs (Tyanova et al., 2016) based on retention time alignment and targeted feature extraction (Weisser and Choudhary, 2017) to integrate respective MS1 areas for all time points and technical replicates.

MS1 areas *x* were normalized to z-scores (standard scores) *z* per MS run by subtracting the mean and dividing by the standard deviation:

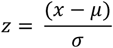

The trajectory of scaled MS1 areas was clustered by k-means unsupervised clustering with 6 seeds using the tslearn (v.0.3.1) python package. All trajectories are related to the first time point by subtracting its median z-score from all other timepoints in the respective analysis.

### Comparison of the HLA-Ligand-Atlas database with IEDB and SysteMHC

All peptides contained in the HLA Ligand Atlas database were compared with peptides listed in the IEDB and SysteMHC databases for HLA-I and HLA-II ligands separately. The list of peptides stored in the IEDB was obtained by downloading the file “epitope_full_v3.zip” from the “Database Export” page. The obtained table was subsequently filtered for positive MS assays, linear peptides and human origin. Peptides with modifications were removed. Peptides stored in the SysteMHC database were obtained by downloading the file “180409_master_final.tgz” from “Builds_for_download” page. The obtained table was subsequently filtered for human as organism.

### Gene ontology (GO)-term enrichment

GO term enrichment analyses were performed with the Panther 15.0 database (Released 2020-02-21) with the integrated “statistical overrepresentation test” (Release 2019-07-11). Gene identifiers of source proteins presented exclusively by either HLA-I or -II allotypes were queried against the “GO cellular component complete” database using the default “Homo sapiens genes” reference list. GO terms were sorted by Fisher’s exact raw p-value, and top 10 scoring terms reported as overrepresented and their corresponding p-values were selected for illustration.

Tissue-specific source proteins were defined as HLA-I or –II source proteins identified exclusively in one tissue across all subjects (Table S5). Gene identifiers of tissue-specific HLA-I and -II source proteins were queried against the “GO biological process complete” database, with the only difference that only the top 5 scoring terms reported as overrepresented were selected for illustration.

### Tissue-specific gene set enrichment

Analogously to the GO-term enrichment, tissue-specific HLA-I and -II source proteins were separately queried against the GTEx database for gene set enrichment analysis. Gene sets with upregulated gene expression profiles per tissue “GTEx_Tissue_Sample_Gene_Expression_Profiles_up” were retrieved using the gseapy implementation (v.0.9.15, 2019-08-07) through the enrichr API. All tissues covered in the HLA Ligand Atlas were matched and compared against all tissues in the GTEx database that co-occur in the HLA Ligand Atlas. Fisher’s exact raw p-values for the enrichment were computed for each pairwise comparison.

### HLA-I and –II peptide yield correlation to expression of immune-related genes

We computed a linear model to compare the median HLA-I peptide yields per tissue with gene expression values (RPKM) of the following genes involved in the HLA-I presentation pathway: HLA-A, HLA-B, HLA-C, immunoproteasome, constitutive proteasome, TAP1, and TAP2. Median HLA-II peptide yields per tissue were correlated to genes involved in the HLA-II presentation pathway: HLA-DRB1, HLA-DRA, HLA-DQB1, HLA-DQA1, HLA-DPB1, HLA-DPA1. The corresponding gene expression values were taken from a previously published study (Boegel et al., 2018).

An ordinary least squares linear model correlating gene expression and log_10_ median HLA-I and -II peptide yields was computed using R (v.3.5) and the corresponding stats (v.3.5) package reporting R^2^, F-statistic p-value, and spearman rho. The cross correlation between all immune related genes and their individual linear models (Figure 3, Figure S5) was computed using R (v.3.5) and the corresponding packages corrplot (v. 0.84) and ggplot2 (v.3.2.1). As the expression levels of the investigated genes are highly covariant (Figure S5A, S5C), the regression would be overfitting when correlating peptide yields to multiple genes involved in the antigen presentation pathway, thus the analysis was limited to a single gene at a time.

### Computation of Jaccard coefficients between samples

We investigated the similarity of immunopeptidomes between tissues and subjects by pairwise comparison of all samples in the HLA Ligand Atlas. Comparisons were performed both on HLA-I and -II level as well as on peptide and source protein level. The Jaccard index was calculated by dividing the set intersection by the set union for all pairwise comparisons:

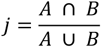

### Identification of cryptic peptides with Peptide-PRISM

Identification of cryptic HLA-I peptides from HLA-I LC-MS/MS data was performed as recently described in detail (Erhard et al., 2020). Briefly, *de novo* peptide sequencing was performed with PEAKS Studio X (Tran et al., 2019; Zhang et al., 2012) (Bioinformatics Solutions Inc., Canada). Top10 sequence candidates were exported for each fragment ion spectrum. Database matching of all sequence candidates and stratified FDR-filtering was performed with Peptide-PRISM using the 6-frame translation of the human genome (HG38) and the 3-frame translation of the human transcriptome (Ensembl 90). Matched peptides were filtered to 10% FDR and peptides were predicted as binder to the corresponding HLA alleles by NetMHCpan-4.0 (Jurtz et al., 2017).

### Retention time model for cryptic peptide validation

Retention time predictions were carried out using the OpenMS (2.5.0) RTModel based on oligo-kernel v-support vector regression (v=0.5, p=0.1, c=1, degree=1, border_length=22, kmer_length=1, Σ=5) (Pfeifer et al., 2007). The model was trained on all peptide identifications of canonical peptides identified with MHCquant and applied to all cryptic peptide identifications resulting from Peptide-PRISM. Predictions were evaluated by applying a linear least square fit to compute the 99% prediction interval around the predicted versus measured retention times using the statsmodels (v.0.11) function wls_prediction_std.

### Synthesis of isotope-labeled peptides

Peptides were synthesized using the Liberty Blue Automated Peptide Synthesizer (CEM) following the standard 9-fluorenylmethyl-oxycarbonyl/*tert*-butyl strategy. After removal from the resin by treatment with trifluoroacetic acid/triisopropylsilane/water (95/2.5/2.5 by vol.) for 1 h, peptides were precipitated from diethyl ether, washed three times with diethyl ether and resuspended in water prior to lyophilization. Purity and identity of the synthesis products were determined by C18-HPLC (Thermo Fisher Scientific, Darmstadt, Germany) and LTQ Orbitrap XL mass spectrometer (Thermo Fisher Scientific), respectively.

### Spectrum validation

We selected 36 cryptic peptides, identified with 1% FDR for spectral validation with isotope-labeled synthetic peptides. Selected peptides were strong binders to the corresponding HLA alleles of the respective subject, with a netMHCpan-4.0 binding rank <0.5.

Isotope-labeled synthetic peptides were spiked into a sample matrix of native HLA eluted peptides from a JY cell line at a concentration of 20 fmol/µl, with the purpose of showing spectrum identity between the native and synthetic peptides.

The spectral similarity *λ* was computed analogous to the normalized spectral contrast angle (Toprak et al., 2014) between eluted peptide spectra and synthetic isotope labeled peptide spectra:

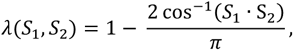

where the spectra were encoded as intensity vectors (*S*_1_and *S*_2_) based on their theoretical *b* and *y* fragment ions by using the mzR (v2.16.2), msdata (v0.20.0) and protViz (v0.4) R packages. Intensities of matching y- and b-ion pairs as encoded in the intensity vectors were compared, thereby avoiding the necessity to correct for the mass shift caused by the isotope label. Peaks present in at least one of the spectra were considered for the cross product (*S*_1_· *S*_2_). Intensities of missing peaks in the one spectrum compared to the other were set to zero.

### Data storage and web interface

Data was stored and managed using the biomedical data-management platform qPortal (Mohr et al., 2018). HLA-I and –II peptides were complemented with their tissue and HLA allotype association and stored in an SQL database. A public web server was implemented that allows users to formulate queries against the database, visualize results and allows data export for further analysis. The web front-end was implemented in HTML, CSS and JavaScript based on the front-end framework Bootstrap 4. The table plugin DataTables was used to provide rapid browsing and filtering for tabular data. Interactive plots were designed using Bokeh and ApexCharts.

## Data availability

The LC-MS/MS immunopeptidomics data comprised in the HLA Ligand Atlas has been deposited to the ProteomeXchange

Consortium via the PRIDE (Perez-Riverol et al., 2019) partner repository with the dataset identifier PXD019643 and the project DOI 10.6019/PXD019643. LC-MS/MS runs and sample not adhering to the implemented quality control thresholds are deposited as well.

The LC-MS/MS immunopeptiodmics data from the three glioblastoma patients can be accessed with the PXD020186, and the project DOI 10.6019/PXD020186.

## Additional resource

The HLA Ligand Atlas: https://hla-ligand-atlas.org/

## REFERENCES

Abelin, J.G., Harjanto, D., Malloy, M., Suri, P., Colson, T., Goulding, S.P., Creech, A.L., Serrano, L.R., Nasir, G., Nasrullah, Y., et al. (2019). Defining HLA-II Ligand Processing and Binding Rules with Mass Spectrometry Enhances Cancer Epitope Prediction. Immunity 51, 766–779.

Almeida, L.G., Sakabe, N.J., de Oliveira, A.R., Silva, M.C.C., Mundstein, A.S., Cohen, T., Chen, Y.T., Chua, R., Gurung, S., Gnjatic, S., et al. (2009). CTdatabase: A knowledge-base of high-throughput and curated data on cancer-testis antigens. Nucleic Acids Res. 37, D816–D819.

Alpízar, A., Marino, F., Ramos-Fernández, A., Lombardía, M., Jeko, A., Pazos, F., Paradela, A., Santiago, C., Heck, A.J.R., and Marcilla, M. (2017). A molecular basis for the presentation of phosphorylated peptides by HLA-B antigens. Mol. Cell. Proteomics 16, 181–193.

Álvaro-Benito, M., Morrison, E., Abualrous, E.T., Kuropka, B., and Freund, C. (2018). Quantification of HLA-DM-Dependent Major Histocompatibility Complex of Class II Immunopeptidomes by the Peptide Landscape Antigenic Epitope Alignment Utility. Front. Immunol. 9, 872.

Aran, D., Camarda, R., Odegaard, J., Paik, H., Oskotsky, B., Krings, G., Goga, A., Sirota, M., and Butte, A.J. (2017). Comprehensive analysis of normal adjacent to tumor transcriptomes. Nat. Commun. 8, 1077.

Ardlie, K.G., DeLuca, D.S., Segrè, A. V., Sullivan, T.J., Young, T.R., Gelfand, E.T., Trowbridge, C.A., Maller, J.B., Tukiainen, T., Lek, M., et al. (2015). The Genotype-Tissue Expression (GTEx) pilot analysis: Multitissue gene regulation in humans. Science (80-.). 348, 648–660.

Barnstable, C.J., Bodmer, W.F., Brown, G., Galfre, G., Milstein, C., Williams, A.F., and Ziegler, A. (1978). Production of monoclonal antibodies to group A erythrocytes, HLA and other human cell surface antigens-new tools for genetic analysis. Cell 14, 9–20.

Bassani-Sternberg, M., Bräunlein, E., Klar, R., Engleitner, T., Sinitcyn, P., Audehm, S., Straub, M., Weber, J., Slotta-Huspenina, J., Specht, K., et al. (2016). Direct identification of clinically relevant neoepitopes presented on native human melanoma tissue by mass spectrometry. Nat. Commun. 7, 13404.

Bassani-Sternberg, M., Chong, C., Guillaume, P., Solleder, M., Pak, H.S., Gannon, P.O., Kandalaft, L.E., Coukos, G., and Gfeller, D. (2017). Deciphering HLA-I motifs across HLA peptidomes improves neo-antigen predictions and identifies allostery regulating HLA specificity. PLoS Comput. Biol. 13, e1005725.

Bichmann, L., Nelde, A., Ghosh, M., Heumos, L., Mohr, C., Peltzer, A., Kuchenbecker, L., Sachsenberg, T., Walz, J.S., Stevanović, S., et al. (2019). MHCquant: Automated and Reproducible Data Analysis for Immunopeptidomics. J. Proteome Res. 18, 3876–3884.

Bilich, T., Nelde, A., Bichmann, L., Roerden, M., Salih, H.R., Kowalewski, D.J., Schuster, H., Tsou, C.C., Marcu, A., Neidert, M.C., et al. (2019). The HLA ligandome landscape of chronic myeloid leukemia delineates novel T-cell epitopes for immunotherapy. Blood 133, 550–565.

Boegel, S., Löwer, M., Bukur, T., Sorn, P., Castle, J.C., and Sahin, U. (2018). HLA and proteasome expression body map. BMC Med. Genomics 11, 36.

Boegel, S., Castle, J.C., Kodysh, J., O’Donnell, T., and Rubinsteyn, A. (2019). Bioinformatic methods for cancer neoantigen prediction. In Progress in Molecular Biology and Translational Science, (Academic Press), pp. 25–60.

Brown, S.D., and Holt, R.A. (2018). Neoantigen characteristics in the context of the complete predicted MHC class I self-immunopeptidome. Oncoimmunology 8, 1556080.

Calis, J.J.A., Maybeno, M., Greenbaum, J.A., Weiskopf, D., De Silva, A.D., Sette, A., Keşmir, C., and Peters, B. (2013). Properties of MHC class I presented peptides that enhance immunogenicity. PLoS Comput. Biol. 9, e1003266–e1003266.

Cameron, B.J., Gerry, A.B., Dukes, J., Harper, J. V., Kannan, V., Bianchi, F.C., Grand, F., Brewer, J.E., Gupta, M., Plesa, G., et al. (2013). Identification of a titin-derived HLA-A1-presented peptide as a cross-reactive target for engineered MAGE A3-directed T cells. Sci. Transl. Med. 5, 197ra103 LP–197ra103.

Caron, E., Aebersold, R., Banaei-Esfahani, A., Chong, C., and Bassani-Sternberg, M. (2017). A Case for a Human Immuno-Peptidome Project Consortium. In Immunity, pp. 203–208.

Chong, C., Müller, M., Pak, H.S., Harnett, D., Huber, F., Grun, D., Leleu, M., Auger, A., Arnaud, M., Stevenson, B.J., et al. (2020). Integrated proteogenomic deep sequencing and analytics accurately identify non-canonical peptides in tumor immunopeptidomes. Nat. Commun. 11, 1293.

Eng, J.K., Hoopmann, M.R., Jahan, T.A., Egertson, J.D., Noble, W.S., and MacCoss, M.J. (2015). A Deeper Look into Comet - Implementation and Features. J. Am. Soc. Mass Spectrom. 26, 1865–1874.

Erhard, F., Dölken, L., Schilling, B., and Schlosser, A. (2020). Identification of the cryptic HLA-I immunopeptidome. Cancer Immunol. Res. canimm.0886.2019.

Ewels, P.A., Peltzer, A., Fillinger, S., Patel, H., Alneberg, J., Wilm, A., Garcia, M.U., Di Tommaso, P., and Nahnsen, S. (2020). The nf-core framework for community-curated bioinformatics pipelines. Nat. Biotechnol. 38, 276–278.

Faridi, P., Purcell, A.W., and Croft, N.P. (2018). In Immunopeptidomics We Need a Sniper Instead of a Shotgun. Proteomics 18, 1700464.

Finotello, F., Rieder, D., Hackl, H., and Trajanoski, Z. (2019). Next-generation computational tools for interrogating cancer immunity. Nat. Rev. Genet. 20, 724–746.

Fortier, M.H., Caron, É., Hardy, M.P., Voisin, G., Lemieux, S., Perreault, C., and Thibault, P. (2008). The MHC class I peptide repertoire is molded by the transcriptome. J. Exp. Med. 205, 595–610.

Freudenmann, L.K., Marcu, A., and Stevanović, S. (2018). Mapping the tumour human leukocyte antigen (HLA) ligandome by mass spectrometry. Immunology 154.

Fritsche, J., Rakitsch, B., Hoffgaard, F., Römer, M., Schuster, H., Kowalewski, D.J., Priemer, M., Stos-Zweifel, V., Hörzer, H., Satelli, A., et al. (2018). Translating Immunopeptidomics to Immunotherapy-Decision-Making for Patient and Personalized Target Selection. Proteomics 18, e1700284–e1700284.

Goldman, J.M., Hibbin, J., Kearney, L., Orchard, K., and Th’ng, K.H. (1982). HLA-DA monoclonal antibodies inhibit the proliferation of normal and chronic granulocytic leukaemia myeloid progenitor cell. Br. J. Haematol. 52, 411–420.

Granados, D.P., Rodenbrock, A., Laverdure, J.-P., Côté, C., Caron-Lizotte, O., Carli, C., Pearson, H., Janelle, V., Durette, C., Bonneil, E., et al. (2016). Proteogenomic-based discovery of minor histocompatibility antigens with suitable features for immunotherapy of hematologic cancers. Leukemia 30, 1344–1354.

Hilf, N., Kuttruff-Coqui, S., Frenzel, K., Bukur, V., Stevanović, S., Gouttefangeas, C., Platten, M., Tabatabai, G., Dutoit, V., van der Burg, S.H., et al. (2019). Actively personalized vaccination trial for newly diagnosed glioblastoma. Nature 565, 240–245.

Iacobuzio-Donahue, C.A., Michael, C., Baez, P., Kappagantula, R., Hooper, J.E., and Hollman, T.J. (2019). Cancer biology as revealed by the research autopsy. Nat. Rev. Cancer 19, 686–697.

Ishii, N., Chiba, M., Iizuka, M., Watanabe, H., Ishioka, T., and Masamune, O. (1992). Expression of MHC class II antigens (HLA-DR, -DP, and -DQ) on human gastric epithelium. Gastroenterol. Jpn. 27, 23–28.

Jiang, L., Wang, M., Lin, S., Jian, R., Li, X., Chan, J., Fang, H., Dong, G., Consortium, Gte., Tang, H., et al. (2019). A Quantitative Proteome Map of the Human Body. BioRxiv 797373.

Jurtz, V., Paul, S., Andreatta, M., Marcatili, P., Peters, B., and Nielsen, M. (2017). NetMHCpan-4.0: Improved Peptide–MHC Class I Interaction Predictions Integrating Eluted Ligand and Peptide Binding Affinity Data. J. Immunol. 199, 3360–3368.

Kaufman, J. (2018). Generalists and Specialists: A New View of How MHC Class I Molecules Fight Infectious Pathogens. Trends Immunol. 39, 367–379.

Kim, M.S., Pinto, S.M., Getnet, D., Nirujogi, R.S., Manda, S.S., Chaerkady, R., Madugundu, A.K., Kelkar, D.S., Isserlin, R., Jain, S., et al. (2014). A draft map of the human proteome. Nature 509, 575–581.

Lander, E.S., Linton, L.M., Birren, B., Nusbaum, C., Zody, M.C., Baldwin, J., Devon, K., Dewar, K., Doyle, M., Fitzhugh, W., et al. (2001). Initial sequencing and analysis of the human genome. Nature 409, 860–921.

Laumont, C.M., Vincent, K., Hesnard, L., Audemard, É., Bonneil, É., Laverdure, J.P., Gendron, P., Courcelles, M., Hardy, M.P., Côté, C., et al. (2018). Noncoding regions are the main source of targetable tumor-specific antigens. Sci. Transl. Med. 10, eaau5516.

Linette, G.P., Stadtmauer, E.A., Maus, M. V, Rapoport, A.P., Levine, B.L., Emery, L., Litzky, L., Bagg, A., Carreno, B.M., Cimino, P.J., et al. (2013). Cardiovascular toxicity and titin cross-reactivity of affinity-enhanced T cells in myeloma and melanoma. Blood 122, 863–871.

Lippolis, J.D., White, F.M., Marto, J.A., Luckey, C.J., Bullock, T.N.J., Shabanowitz, J., Hunt, D.F., and Engelhard, V.H. (2002). Analysis of MHC Class II Antigen Processing by Quantitation of Peptides that Constitute Nested Sets. J. Immunol. 169, 5089 LP – 5097.

Löffler, M.W., Mohr, C., Bichmann, L., Freudenmann, L.K., Walzer, M., Schroeder, C.M., Trautwein, N., Hilke, F.J., Zinser, R.S., Mühlenbruch, L., et al. (2019). Multi-omics discovery of exome-derived neoantigens in hepatocellular carcinoma. Genome Med. 11, 28.

Di Marco, M., Schuster, H., Backert, L., Ghosh, M., Rammensee, H.-G., and Stevanović, S. (2017). Unveiling the Peptide Motifs of HLA-C and HLA-G from Naturally Presented Peptides and Generation of Binding Prediction Matrices. J. Immunol. 199, 2639–2651.

Marino, F. (2018). Gaining Insight Into Posttranslationally Modified HIV Antigens and Their Underlying Characteristics. Proteomics 18, 1800041.

Melé, M., Ferreira, P.G., Reverter, F., DeLuca, D.S., Monlong, J., Sammeth, M., Young, T.R., Goldmann, J.M., Pervouchine, D.D., Sullivan, T.J., et al. (2015). The human transcriptome across tissues and individuals. Science (80-.). 348, 660–665.

Mohr, C., Friedrich, A., Wojnar, D., Kenar, E., Polatkan, A.C., Codrea, M.C., Czemmel, S., Kohlbacher, O., and Nahnsen, S. (2018). qPortal: A platform for data-driven biomedical research. PLoS One 13, e0191603.

Müller, M., Gfeller, D., Coukos, G., and Bassani-Sternberg, M. (2017). “Hotspots” of antigen presentation revealed by human leukocyte antigen ligandomics for neoantigen prioritization. Front. Immunol. 8, 1367.

Mylonas, R., Beer, I., Iseli, C., Chong, C., Pak, H.-S., Gfeller, D., Coukos, G., Xenarios, I., Müller, M., and Bassani-Sternberg, M. (2018). Estimating the Contribution of Proteasomal Spliced Peptides to the HLA-I Ligandome*. Mol. & Cell. Proteomics 17, 2347 LP – 2357.

Ott, P.A., Hu, Z., Keskin, D.B., Shukla, S.A., Sun, J., Bozym, D.J., Zhang, W., Luoma, A., Giobbie-Hurder, A., Peter, L., et al. (2017). An immunogenic personal neoantigen vaccine for patients with melanoma. Nature.

Ouspenskaia, T., Law, T., Clauser, K.R., Klaeger, S., Sarkizova, S., Aguet, F., Li, B., Christian, E., Knisbacher, B.A., Le, P.M., et al. (2020). Thousands of novel unannotated proteins expand the MHC I immunopeptidome in cancer. BioRxiv 2020.02.12.945840.

Pawelec, G., Ziegler, A., and Wernet, P. (1985). Dissection of human allostimulatory determinants with cloned T cells: Stimulation inhibition by monoclonal antibodies TÜ22, 34, 36, 37, 39, 43, and 58 against distinct human MHC class II molecules. Hum. Immunol. 12, 165–176.

Perez-Riverol, Y., Csordas, A., Bai, J., Bernal-Llinares, M., Hewapathirana, S., Kundu, D.J., Inuganti, A., Griss, J., Mayer, G., Eisenacher, M., et al. (2019). The PRIDE database and related tools and resources in 2019: improving support for quantification data. Nucleic Acids Res. 47, D442–D450.

Pfeifer, N., Leinenbach, A., Huber, C.G., and Kohlbacher, O. (2007). Statistical learning of peptide retention behavior in chromatographic separations: a new kernel-based approach for computational proteomics. BMC Bioinformatics 8, 468.

Racle, J., Michaux, J., Rockinger, G.A., Arnaud, M., Bobisse, S., Chong, C., Guillaume, P., Coukos, G., Harari, A., Jandus, C., et al. (2019). Robust prediction of HLA class II epitopes by deep motif deconvolution of immunopeptidomes. Nat. Biotechnol. 37, 1283–1286.

Rammensee, H.-G. (1995). Chemistry of peptides associated with MHC class I and class II molecules. Curr. Opin. Immunol. 7, 85–96.

Rammensee, H.-G., Rötzschke, O., and Falk, K. (1993a). MHC Class I-Restricted Antigen Processing — Lessons from Natural Ligands. In Chemical Immunology and Allergy, pp. 113–133.

Rammensee, H.-G., Bachmann, J., Emmerich, N.P.N., Bachor, O.A., and Stevanović, S. (1999). SYFPEITHI: database for MHC ligands and peptide motifs. Immunogenetics 50, 213–219.

Rammensee, H.G., Rötzschke, O., and Falk, K. (1993b). Self tolerance of natural MHC class I ligands. Int. Rev. Immunol. 10, 291–300.

Reiter, L., Claassen, M., Schrimpf, S.P., Jovanovic, M., Schmidt, A., Buhmann, J.M., Hengartner, M.O., and Aebersold, R. (2009). Protein identification false discovery rates for very large proteomics data sets generated by tandem mass spectrometry. Mol. Cell. Proteomics 8, 2405–2417.

Reynisson, B., Barra, C., Kaabinejadian, S., Hildebrand, W.H., Peters, B., and Nielsen, M. (2020). Improved prediction of MHC II antigen presentation through integration and motif deconvolution of mass spectrometry MHC eluted ligand data. J. Proteome Res. 1535–3893.

Rolfs, Z., Solntsev, S.K., Shortreed, M.R., Frey, B.L., and Smith, L.M. (2019). Global Identification of Post-Translationally Spliced Peptides with Neo-Fusion. J. Proteome Res. 18, 349–358.

Röst, H.L., Sachsenberg, T., Aiche, S., Bielow, C., Weisser, H., Aicheler, F., Andreotti, S., Ehrlich, H.C., Gutenbrunner, P., Kenar, E., et al. (2016). OpenMS: A flexible open-source software platform for mass spectrometry data analysis. Nat. Methods 13, 741–748.

Sahin, U., Derhovanessian, E., Miller, M., Kloke, B.P., Simon, P., Löwer, M., Bukur, V., Tadmor, A.D., Luxemburger, U., Schrörs, B., et al. (2017). Personalized RNA mutanome vaccines mobilize poly-specific therapeutic immunity against cancer. Nature 547, 222–226.

Savitski, M.M., Wilhelm, M., Hahne, H., Kuster, B., and Bantscheff, M. (2015). A Scalable Approach for Protein False Discovery Rate Estimation in Large Proteomic Data Sets. Mol. Cell. Proteomics 14, 2394–2404.

Schuster, H., Peper, J.K., Bösmüller, H.C., Röhle, K., Backert, L., Bilich, T., Ney, B., Löffler, M.W., Kowalewski, D.J., Trautwein, N., et al. (2017). The immunopeptidomic landscape of ovarian carcinomas. Proc. Natl. Acad. Sci. U. S. A. 114, E9942–E9951.

Shao, W., Pedrioli, P.G.A., Wolski, W., Scurtescu, C., Schmid, E., Vizcaíno, J.A., Courcelles, M., Schuster, H., Kowalewski, D., Marino, F., et al. (2018). The SysteMHC Atlas project. Nucleic Acids Res. 46, D1237–D1247.

Stern, L.J., Brown, J.H., Jardetzky, T.S., Gorga, J.C., Urban, R.G., Strominger, J.L., and Wiley, D.C. (1994). Crystal structure of the human class II MHC protein HLA-DR1 complexed with an influenza virus peptide. Nature 368, 215–221.

Szolek, A., Schubert, B., Mohr, C., Sturm, M., Feldhahn, M., and Kohlbacher, O. (2014). OptiType: precision HLA typing from next-generation sequencing data. Bioinformatics 30, 3310–3316.

The, M., MacCoss, M.J., Noble, W.S., and Käll, L. (2016). Fast and Accurate Protein False Discovery Rates on Large-Scale Proteomics Data Sets with Percolator 3.0. J. Am. Soc. Mass Spectrom. 27, 1719–1727.

Toprak, U.H., Gillet, L.C., Maiolica, A., Navarro, P., Leitner, A., and Aebersold, R. (2014). Conserved peptide fragmentation as a benchmarking tool for mass spectrometers and a discriminating feature for targeted proteomics. Mol. Cell. Proteomics 13, 2056–2071.

Tran, N.H., Qiao, R., Xin, L., Chen, X., Liu, C., Zhang, X., Shan, B., Ghodsi, A., and Li, M. (2019). Deep learning enables de novo peptide sequencing from data-independent-acquisition mass spectrometry. Nat. Methods 16, 63–66.

Tyanova, S., Temu, T., and Cox, J. (2016). The MaxQuant computational platform for mass spectrometry-based shotgun proteomics. Nat. Protoc. 11, 2301–2319.

Uhlén, M., Fagerberg, L., Hallström, B.M., Lindskog, C., Oksvold, P., Mardinoglu, A., Sivertsson, Å., Kampf, C., Sjöstedt, E., Asplund, A., et al. (2015). Tissue-based map of the human proteome. Science (80-.). 347, 1260419.

Venter, C.J., Adams, M.D., Myers, E.W., Li, P.W., Mural, R.J., Sutton, G.G., Smith, H.O., Yandell, M., Evans, C.A., Holt, R.A., et al. (2001). The sequence of the human genome. Science (80-.). 291, 1304–1351.

Vita, R., Overton, J.A., Greenbaum, J.A., Ponomarenko, J., Clark, J.D., Cantrell, J.R., Wheeler, D.K., Gabbard, J.L., Hix, D., Sette, A., et al. (2015). The immune epitope database (IEDB) 3.0. Nucleic Acids Res. 43, D405–D412.

Vizcaíno, J.A., Kubiniok, P., Kovalchik, K.A., Ma, Q., Duquette, J.D., Mongrain, I., Deutsch, E.W., Peters, B., Sette, A., Sirois, I., et al. (2020). The Human Immunopeptidome Project: A Roadmap to Predict and Treat Immune Diseases. Mol. & Cell. Proteomics 19, 31 LP – 49.

Wang, C., Gu, Y., Zhang, K., Xie, K., Zhu, M., Dai, N., Jiang, Y., Guo, X., Liu, M., Dai, J., et al. (2016). Systematic identification of genes with a cancer-testis expression pattern in 19 cancer types. Nat. Commun. 7, 10499.

Wang, D., Eraslan, B., Wieland, T., Hallström, B., Hopf, T., Zolg, D.P., Zecha, J., Asplund, A., Li, L., Meng, C., et al. (2019). A deep proteome and transcriptome abundance atlas of 29 healthy human tissues. Mol. Syst. Biol. 15, e8503.

Weinzierl, A.O., Lemmel, C., Schoor, O., Müller, M., Krüger, T., Wernet, D., Hennenlotter, J., Stenzl, A., Klingel, K., Rammensee, H.G., et al. (2007). Distorted relation between mRNA copy number and corresponding major histocompatibility complex ligand density on the cell surface. Mol. Cell. Proteomics 6, 102–113.

Weisser, H., and Choudhary, J.S. (2017). Targeted Feature Detection for Data-Dependent Shotgun Proteomics. J. Proteome Res. 16, 2964–2974.

Wilhelm, M., Schlegl, J., Hahne, H., Gholami, A.M., Lieberenz, M., Savitski, M.M., Ziegler, E., Butzmann, L., Gessulat, S., Marx, H., et al. (2014). Mass-spectrometry-based draft of the human proteome. Nature 509, 582–587.

Zhang, J., Xin, L., Shan, B., Chen, W., Xie, M., Yuen, D., Zhang, W., Zhang, Z., Lajoie, G.A., and Ma, B. (2012). PEAKS DB: De Novo Sequencing Assisted Database Search for Sensitive and Accurate Peptide Identification. Mol. Cell. Proteomics 11, M111.010587.

