## Supplemental Figures for "The HLA Ligand Atlas - A resource of natural HLA ligands presented on benign tissues"

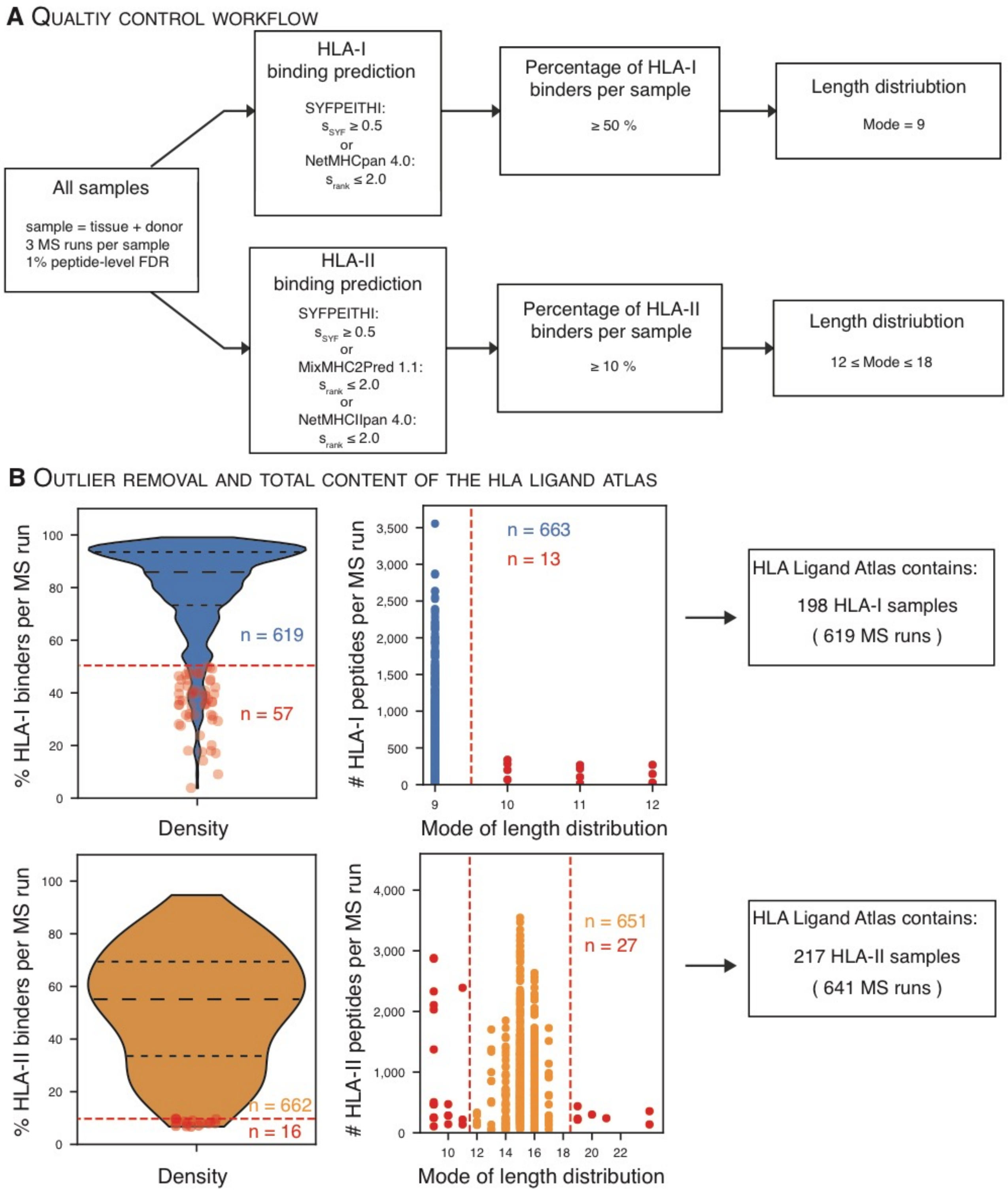
A Qualtiy control workflow
All samples
sample = tissue + donor
3 MS runs per sample
1% peptide-level FDR
HLA-I
binding prediction
SYFPEITHI:
s_SYF ≥ 0.5
or
NetMHCpan 4.0:
s_rank ≤ 2.0
Percentage of HLA-I binders per sample
≥ 50 %
Length distriubtion
Mode = 9
HLA-II
binding prediction
SYFPEITHI:
s_SYF ≥ 0.5
or
MixMHC2Pred 1.1:
s_rank ≤ 2.0
or
NetMHCIIpan 4.0:
s_rank ≤ 2.0
Percentage of HLA-II binders per sample
≥ 10 %
Length distriubtion
12 ≤ Mode ≤ 18
B Outlier removal and total content of the HLA ligand atlas
% HLA-I binders per MS run
n = 619
n = 57
Density
### HLA-I peptides per MS run
n = 663
n = 13
Mode of length distribution
HLA Ligand Atlas contains:
198 HLA-I samples
( 619 MS runs )
% HLA-II binders per MS run
n = 662
n = 16
Density
### HLA-II peptides per MS run
n = 651
n = 27
Mode of length distribution
HLA Ligand Atlas contains:
217 HLA-II samples
( 641 MS runs )

**Supplemental Figure S1: Computational workflow and quality control thresholds (related to Figure 1).**

(A) Data from three LC-MS/MS runs (technical replicates) per sample (one tissue from one subject) were processed with MHCquant with a local peptide-level FDR of 1%. Identified peptides were categorized into peptides predicted as strong, weak, or non-binders. HLA-I peptides were predicted by SYFPEITHI and NetMHCpan 4.0, employing a cut-off threshold of $s_{\mathrm{SYF}} \geq 0.5$ and $s_{\mathrm{rank}} \leq 2.0$, respectively. HLA-II peptides were predicted with MixMHCPred, NetMHCIIpan 4.0 and SYFPEITHI with equal thresholds. Samples with less than 50% and 10% predicted binders per technical replicate for HLA-I and HLA-II immunopeptidomes were excluded from further analysis and are not included in the HLA Ligand Atlas database release. Finally, we employed a stringent cut-off concerning the mode of the peptide length distribution for HLA-I (mode equal to 9) and -II (mode between 12 to18) immunopeptidomes.

(B) LC-MS/MS runs not pertaining to the aforementioned QC thresholds (dashed red lines) concerning the percentage of predicted HLA-binders and the peptide length distribution were not included in the HLA Ligand Atlas database release: Violin plots (left) depict the percentage of peptides predicted to be HLA-binders per LC-MS/MS run and the quality control cut-off for LC-MS/MS runs employed for HLA-I and -II immunopeptidomes. Dot plots (center) depict the mode of the peptide length distribution encountered per LC-MS/MS run and the quality control cut-off employed for HLA-I and -II. The number of LC-MS/MS runs (HLA-I - blue and HLA-II orange) failing the QC thresholds is indicated by red dots. The final number of LC-MS/MS runs and samples selected for inclusion into the HLA Ligand Atlas database release and data analysis are indicated on the right and differ for HLA-I and –II experiments.

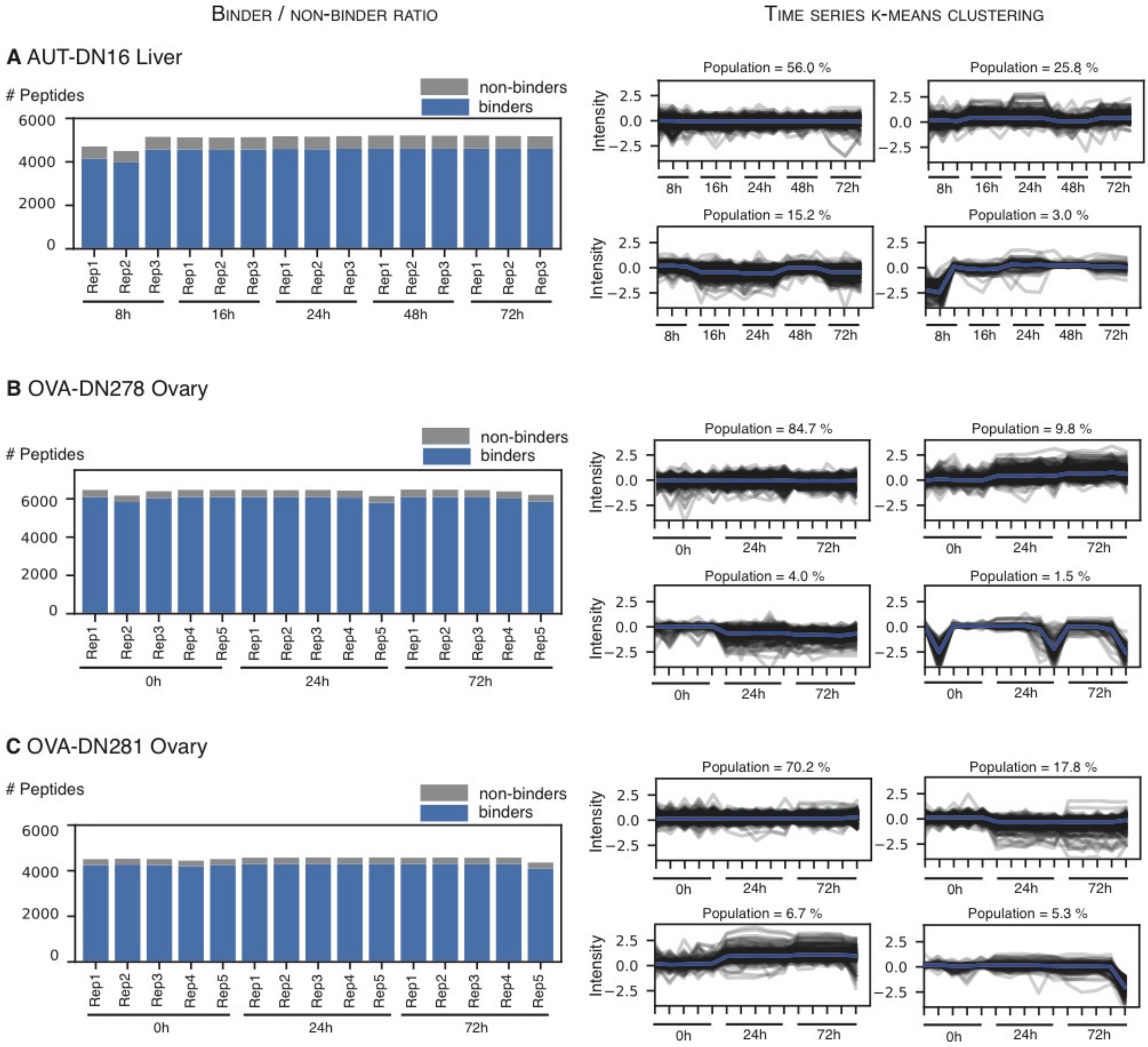


**Supplemental Figure S2: Time-Series Experiment (related to Figure 1).**

The time series experiment was carried out on three biological samples from three subjects (A: AUT-DN16 Liver, B: OVA-DN278, C: OVA-DN281).

A-C: Bar plots (left hand side) indicate the number of identified HLA-I predicted binders (blue) and predicted non-binders (grey) across technical replicates and timepoints for each timeseries.

Time series (right hand side) indicate individual clusters (K-Means using four seeds) of trajectories of quantified MS1 intensity across technical replicates and timepoints. Intermediate trend lines for each cluster are indicated in blue and the percentage of trajectories populating a given cluster is annotated at the top edge of each plot. The trajectories of all timepoints were set in relation to the initial timepoint. The analysis reveals that the number of identified peptides and their percentage of predicted HLA-binders is constant and that most trajectories do not vary much across time points.

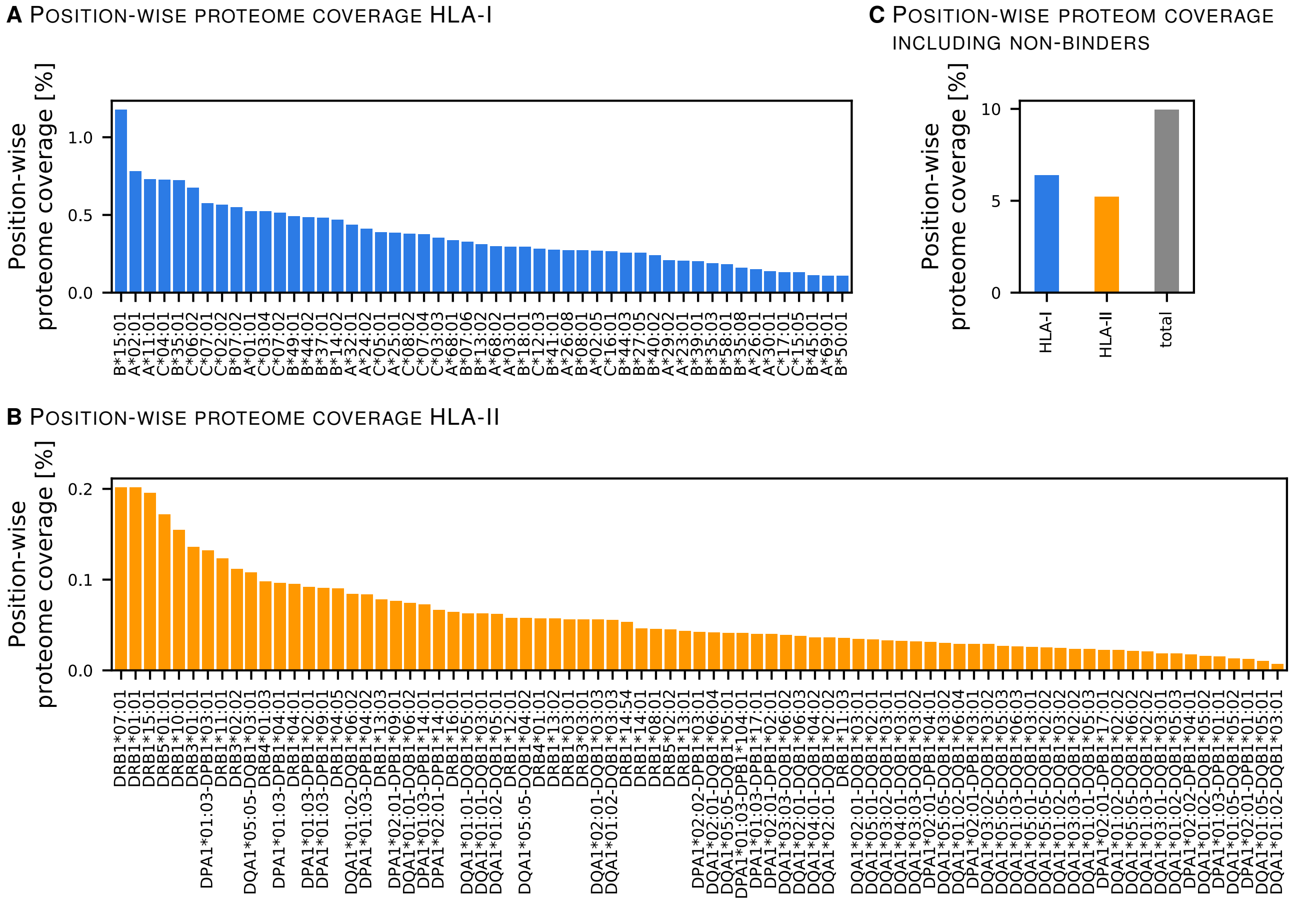


**Supplemental Figure S3: Position-wise proteome coverage (related to Figure 2).**

(A, B) Analysis of the different propensities of HLA-I (blue) and –II (orange), allotypes concerning the presentation of peptides. In turn different percentages of the source proteome are position-wise covered. For example, HLA-B*15:01 ligands cover over 1% of the HLA-I source proteome.

(C) Global analysis of the position-wise coverage of all proteins presented on either HLA-I (blue), HLA-II (orange), or both independent of specific allotypes.

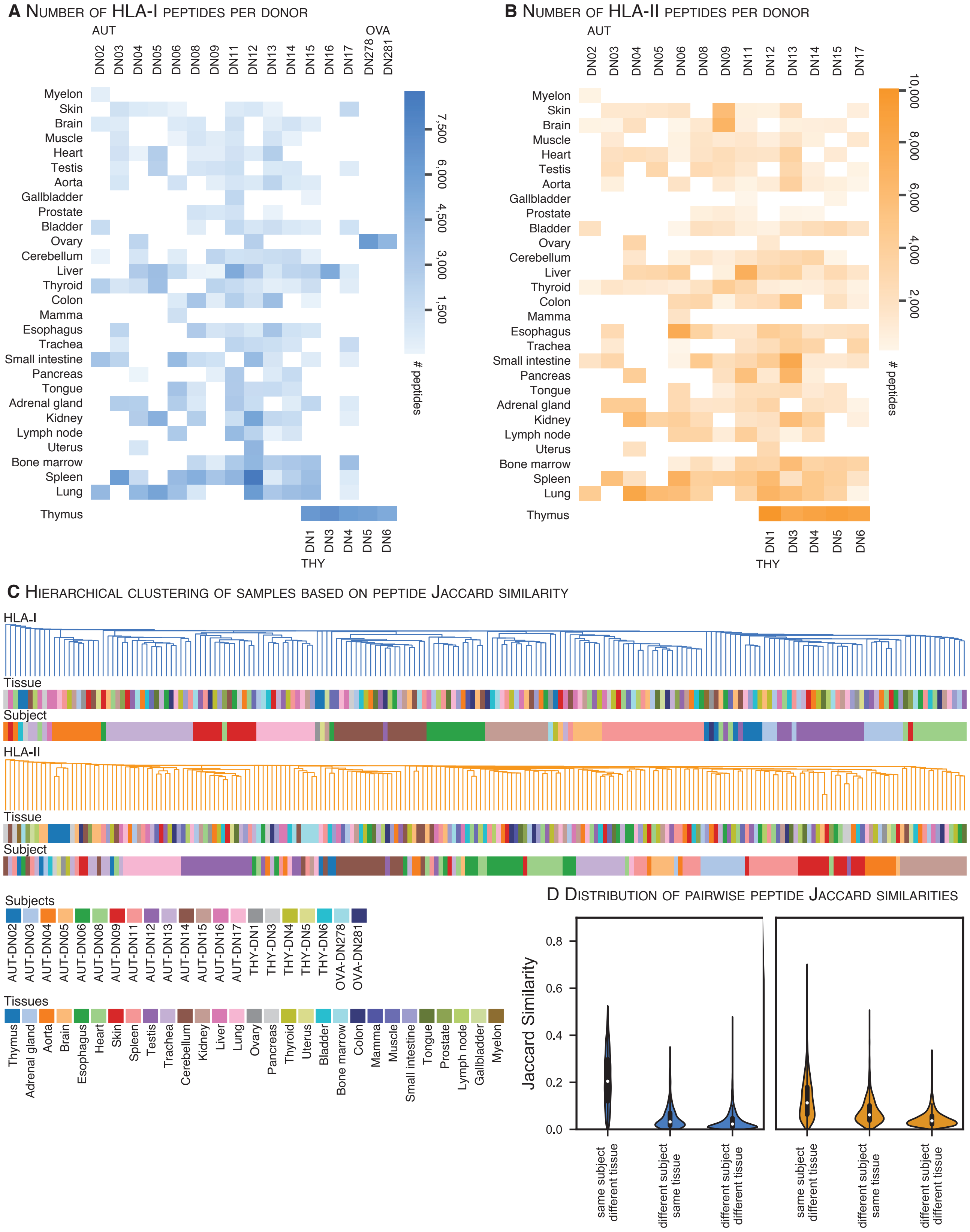
A Number of HLA-I peptides per donor
AUT
OVA
DN02 DN03 DN04 DN05 DN06 DN08 DN09 DN11 DN12 DN13 DN14 DN15 DN16 DN17 DN278 DN281
Myelon
Skin
Brain
Muscle
Heart
Testis
Aorta
Gallbladder
Prostate
Bladder
Ovary
Cerebellum
Liver
Thyroid
Colon
Mamma
Esophagus
Trachea
Small intestine
Pancreas
Tongue
Adrenal gland
Kidney
Lymph node
Uterus
Bone marrow
Spleen
Lung
Thymus
DN1 DN3 DN4 DN5 DN6
THY
7,500
6,000
4,500
3,000
1,500
### peptides
B Number of HLA-II peptides per donor
AUT
DN02 DN03 DN04 DN05 DN06 DN08 DN09 DN11 DN12 DN13 DN14 DN15 DN17
Myelon
Skin
Brain
Muscle
Heart
Testis
Aorta
Gallbladder
Prostate
Bladder
Ovary
Cerebellum
Liver
Thyroid
Colon
Mamma
Esophagus
Trachea
Small intestine
Pancreas
Tongue
Adrenal gland
Kidney
Lymph node
Uterus
Bone marrow
Spleen
Lung
Thymus
DN1 DN3 DN4 DN5 DN6
THY
10,000
8,000
6,000
4,000
2,000
### peptides
C Hierarchical clustering of samples based on peptide Jaccard similarity
HLA-I
Tissue
Subject
HLA-II
Tissue
Subject
Subjects
AUT-DN02 AUT-DN03 AUT-DN04 AUT-DN05 AUT-DN06 AUT-DN08 AUT-DN09 AUT-DN11 AUT-DN12 AUT-DN13 AUT-DN14 AUT-DN15 AUT-DN16 AUT-DN17 THY-DN1 THY-DN3 THY-DN4 THY-DN5 THY-DN6 OVA-DN278 OVA-DN281
Tissues
Thymus Adrenal gland Aorta Brain Esophagus Heart Skin Spleen Testis Trachea Cerebellum Kidney Liver Lung Ovary Pancreas Thyroid Uterus Bladder Bone marrow Colon Mamma Muscle Small intestine Tongue Prostate Lymph node Gallbladder Myelon
D Distribution of pairwise peptide Jaccard similarities
Jaccard Similarity
0.0
0.2
0.4
0.6
0.8
same subject different tissue
different subject same tissue
different subject different tissue
same subject different tissue
different subject same tissue
different subject different tissue

**Supplemental Figure S4: Distribution of peptide identifications and yields across tissues and subjects and peptide-based hierarchical sample clustering (related to Figure 2D, E, and Figure 4A).**

(A, B) HLA-I peptide yields per tissue and subject are illustrated in a heatmap for HLA-I and –II. The color range is in accordance with the number of peptides identified in each sample as indicated in the legend on the right (HLA-I – blue, HLA-II – orange).

(C) Pairwise hierarchical clustering of samples based on the Jaccard similarity between HLA-I (blue) and HLA-II (orange) peptides. The dendrogram illustrates the nearest neighbor based on the similarity between tissues and subjects. See Figure 2D.

(D) Violin plots illustrate the distribution of the peptide Jaccard similarity index for each pairwise comparison between the same subject - different tissues; different subjects - the same tissue, and different subject - different tissues. Results confirmed for HLA-I and -II source proteins Figure 2D and E, respectively.

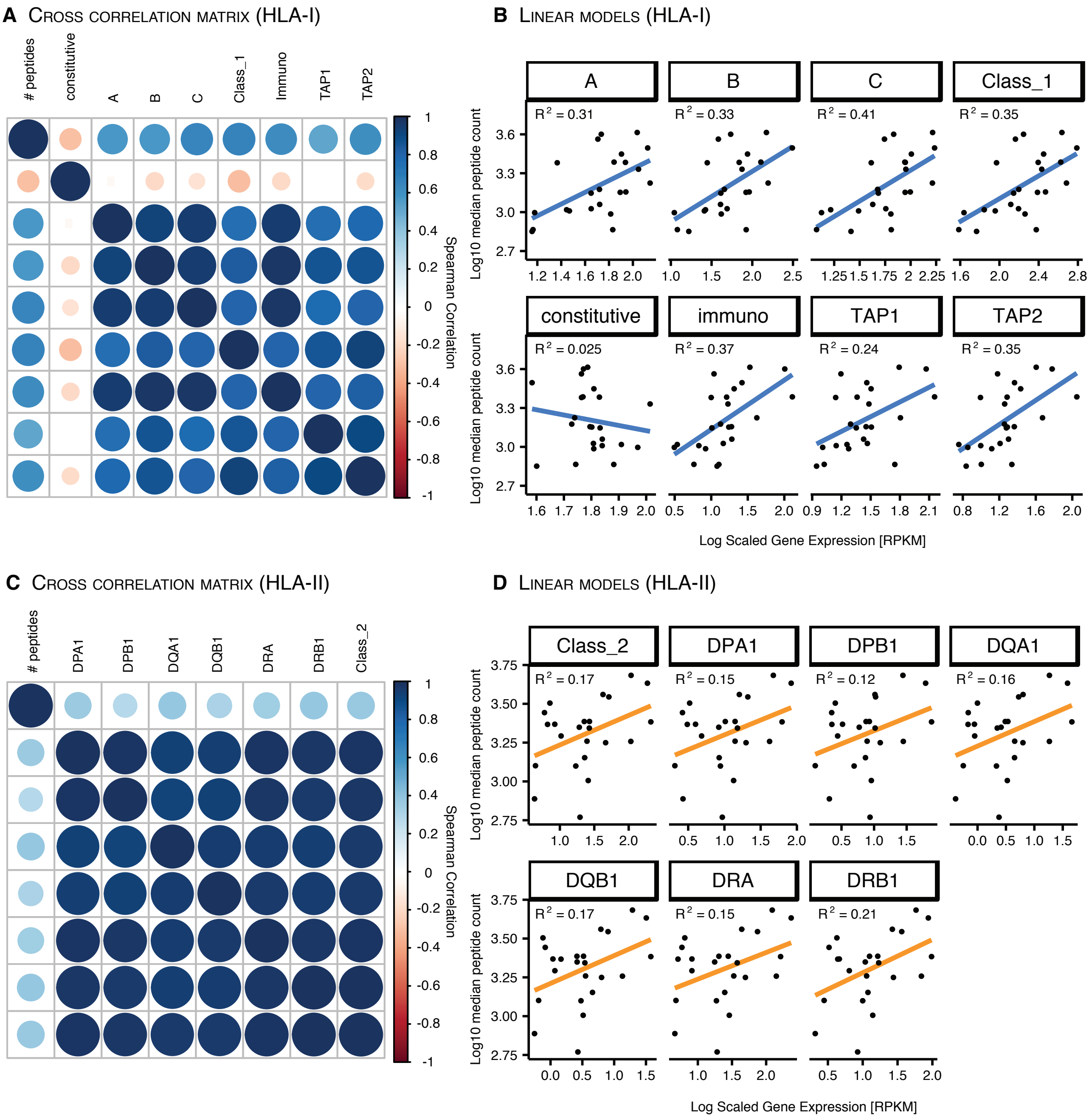


**Supplemental Figure 5: Linear models of gene expression across tissues (related to Figure 3).**

The correlation of HLA-I and -II peptide yields across tissues was assessed with respect to the expression levels (RPKM) of multiple immune related genes of the HLA-I and -II antigen presentation pathway. (RNA expression values were taken from a different publication (Boegel et al., 2018)

A, C) Symmetrical cross-correlation matrices were generated, illustrating the spearman correlation coefficients (rho) of the number of identified HLA-I and -II ligands with expression levels (RPKM) of relevant genes in the HLA-I and -II antigen presentation pathways and among each other. The color-coded dots and their size represent the degree (Spearman rho) of positive (blue) or negative correlation (red).

B, D) In addition, linear models illustrate HLA-I and -II immunopeptidome yields correlated to log scaled gene expression values (RPKM) of genes involved in the HLA-I and -II antigen processing pathway, respectively. The $R^2$ correlation coefficient is depicted in the top left corner for each comparison.

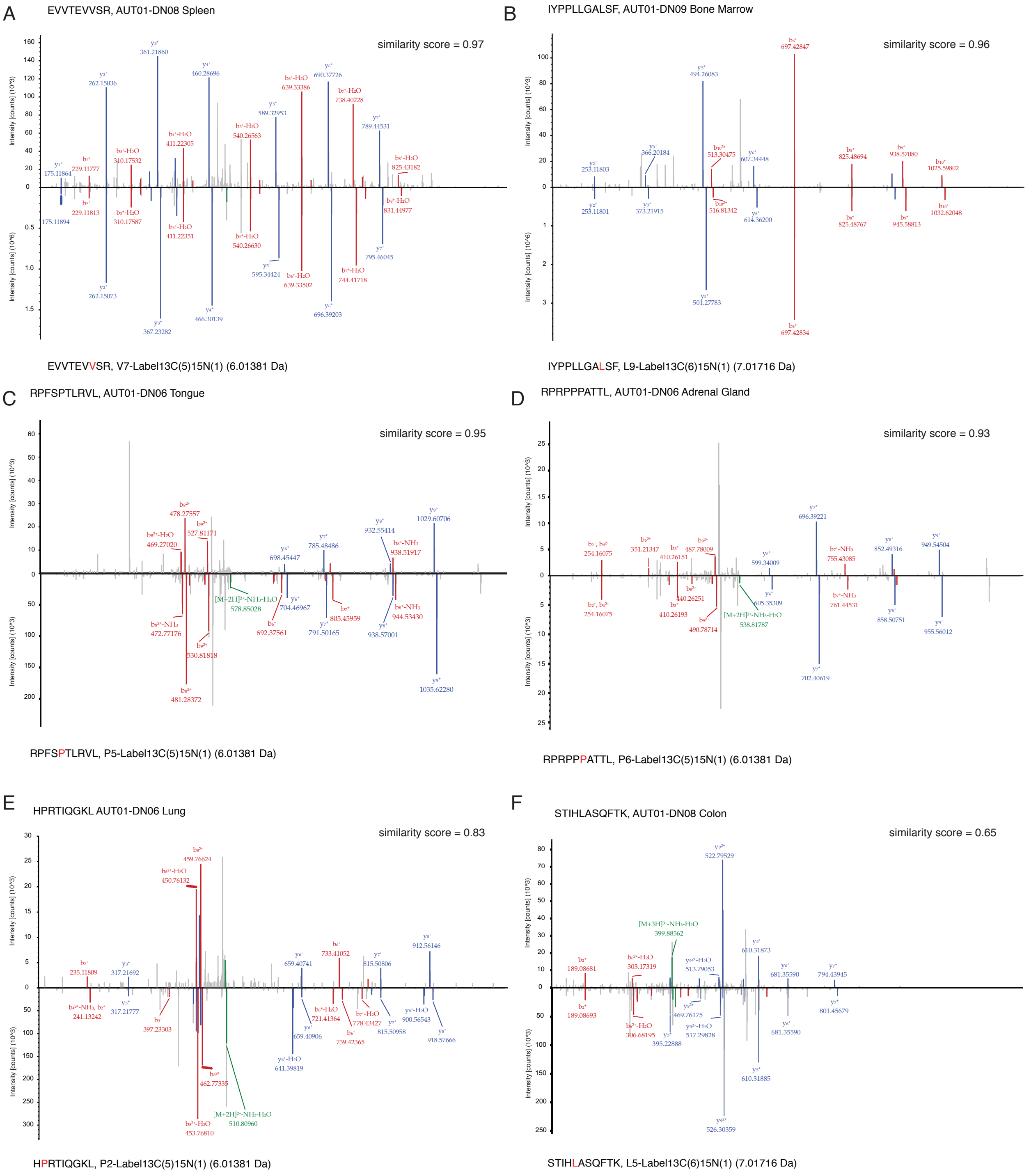


**Supplemental Figure 6: Manual validation of cryptic peptide identifications with isotope labeled synthetic peptides (related to Figure 6).**

A-F) Six exemplary spectral comparisons depict cryptic peptides identified in diverse tissues and subjects (upper spectrum) related to their synthetic isotope labeled counterpart (lower spectrum). The isotope labeled amino acid is highlighted in red and its corresponding label and mass are annotated. Matching b- (red), y-ions (blue) and neutral losses (green) and their corresponding fragment masses are annotated to each peak in the spectra. The spectral similarity score of the given comparison is annotated in the top right.
